# Joint Variable Selection for Omic Biomarkers in Time-to-Event Data

**DOI:** 10.64898/2026.04.30.721585

**Authors:** Jakub Bajzik, Al Depope, Yasaman Zolfimoselo, Alexander Sharipov, Alexandra Lesayova, Holger Klein, Anne Richmond, Spyros Vernardis, Arturas Grauslys, Sergej Andrejev, Aleksej Zelezniak, Markus Ralser, Riccardo E. Marioni, Marco Mondelli, Matthew R. Robinson

## Abstract

The incidence of the vast majority of neurodegenerative, cancer, and metabolic diseases generally increases exponentially with age. In large-scale biobanks, linking time-to-diagnosis information in electronic health records to multiple genomic (“multiomics”) measures has the potential to reveal the genes and biological pathways involved in the disease onset and progression. To date, association testing has commonly been conducted by testing one variable at a time using semiparametric Cox proportional hazards (CoxPH) models, which ignores correlation structure and increases the risk of false discoveries. To address these issues, we introduce a novel fully parametric Bayesian computational method, vampW, based on the Vector Approximate Message Passing framework applied to a Weibull model. vampW jointly models correlated features, while providing an interpretable hazard structure, producing a continuous survival curve, and incorporating prior knowledge. In an extensive simulation study, we demonstrate that joint modeling of omics data and time-to-event outcomes with vampW, substantially reduces false discoveries in comparison to marginal testing and other forms of joint CoxPH models. In 53,018 individuals from the UK Biobank, vampW identifies 219 protein associations with 24 disease outcomes, most of which are not among the top marginal discoveries. We further correct protein levels for exponential age effects, identifying 1,308 associations and highlighting the sensitivity of the analysis to age-correction methodology. Our findings replicate in independent cohorts using different measurement technologies, within data from Iceland and a novel Generation Scotland proteomics dataset. vampW also achieves significant improvement in the prediction of disease onset times: across 14 outcomes, it reduces the root mean squared error by over 30% and 26%, when compared to CoxPH variants and the deep learning approach DeepSurv, respectively, while maintaining predictive utility in minority populations. In summary, vampW offers accurate and interpretable variable selection and out-of-sample prediction within a single computational framework, making it a powerful tool for dissecting the genomic architecture of common complex disease onset.

## Introduction

Aging is the primary risk factor for the vast majority of common chronic, noncommunicable diseases that cause illness and death worldwide. The relationship is generally exponential, where disease risk and incidence increase sharply with advancing age [1]. The potential genes and pathways involved in disease onset and progression can be identified by associating time-to-diagnosis information contained within electronic health records to multiple genomic (“multiomics”) measures [2, 3]. The prognostic biomarkers identified can then provide predictors of age at disease onset, facilitating patient screening programs and preventative medicine [2, 3].

To date, the two main tasks in time-to-event modeling of multiomics data—association testing and out-of-sample prediction of disease onset age—are almost always conducted using semiparametric Cox proportional hazards (PH) models. More precisely, association testing of disease onset age in genomic studies has predominantly been conducted with marginal CoxPH testing [2, 4, 5]. This corresponds to analyzing one genomic feature at a time and it ignores pervasive correlations among genomic measures [6], making it difficult to identify which proteins are most likely driving the observed patterns, with an increased risk of false discoveries [7]. For prediction tasks, penalized CoxPH models [8] are frequently employed [2, 9], enabling the simultaneous modeling of many markers while shrinking or zeroing out a subset of coefficients. Cross-validation is commonly used to select the optimal level of penalization and stability selection [10] can further be applied to reduce the impact of stochastic variability in model fitting when identifying associated predictors. In practice, this involves fitting multiple models with fixed hyperparameters and selecting only stable features—those that are included in a significant proportion of the models. The CoxPH model relies on a partial likelihood that ignores the baseline hazard and the exact spacing between events. While this makes semi-parametric Cox regression robust and widely applicable, it is statistically less efficient because it depends only on the ordering of event times rather than their absolute values. Hence, it does not provide direct predictions of absolute event times. Instead, it yields relative risk scores with respect to other individuals in the dataset.

Here, we provide an alternative to semi-parametric models by introducing vampW, a joint estimation procedure based on Vector Approximate Message Passing (VAMP) [11, 12] that employs a fully parametric Weibull survival model. The methodological novelty of vampW lies in the introduction of Weibull channel denoisers and in handling right-censoring. Using large-scale proteomics data linked to electronic health records [3, 13, 14], we conduct extensive simulations and benchmarking analyses to demonstrate that joint modeling of time-to-event outcomes using vampW substantially reduces false-positive protein associations, as compared to conventional one-protein-at-a-time testing and other forms of CoxPH models. Applying vampW to 24 disease outcomes in the UK Biobank, we identify 1,308 and 219 protein–disease associations in models with and without exponential age correction, respectively, highlighting the sensitivity of protein–outcome associations to the choice of age-correction method. In addition to identifying several well-established biomarkers, such as *APOE* with Alzheimer, *REN* with hypertension and *KLK3* with prostate cancer, we find novel associations that may point to previously unrecognized disease pathways. In particular, we identify *CXCL17*, previously reported as a potential biomarker for colorectal cancer and ischemic heart disease, as being associated with depression. The majority of vampW findings replicate in an independent Icelandic cohort, when adjusted for overlap in proteins and outcomes, despite different measurement technologies between the studies. We also analyze novel Generation Scotland proteomics data generated using mass-spectrometry technology, which further supports our findings. Moreover, across 14 outcomes with sufficient prevalence for out-of-sample evaluation, vampW predicts age at onset with an average root mean squared error which is over 30% and 26% lower than CoxPH variants and the deep learning approach DeepSurv [15], respectively. We then show that the polygenic risk scores generated by vampW are transferable across diverse genetic ancestries, even in minority populations. Finally, we show that vampW outperforms deep learning methods in the data-scarce regimes on standard survival benchmarking datasets, demonstrating the utility of our approach across datasets in the life sciences.

## Results

### Overview of the approach

Our aim is to jointly model the data in a design matrix ***X*** ∈ ℝ^*N* ×*P*^, containing observations on *N* individuals and *P* biological markers, ensuring that we control for the correlation structure. We consider a Weibull model for time-to-event observations, **y** = [*y*_1_, …, *y*_*N*_]^⊤^, where *y*_*i*_ denotes the time-to-event (diagnosis) for individual *i*. The model is formulated as follows:

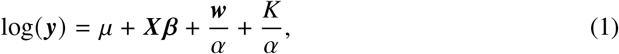

where ***⊤*** = [*w*_1_, …, *w*_*N*_] is a random vector containing independent, Gumbel (0, 1) distributed variables, *α* is the Weibull shape parameter, ***β*** is the vector of marker effects, *μ* is the model intercept, and *K* is the Euler-Mascheroni constant. We normalize the entries of ***X*** to have zero mean and unit variance across columns. In order to allow for a range of effect magnitudes, we select the prior on ***β*** to be an adaptive spike-and-slab form:

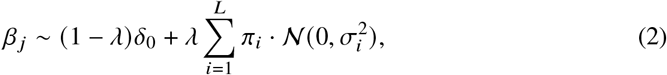

where *L* denotes number of mixture components, (*π*_1_, …, *π*_*L*_) are inclusion probabilities, 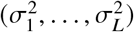 are mixture-specific variances, *δ*_0_ is a Dirac spike function at 0, and *λ* is the sparsity rate (for full details see Methods).

To fit the parameters of the model outlined in Equation 1, we introduce a novel generalized vector approximate message passing algorithm (called *vampW*), which adaptively estimates the prior-specific parameters from Equation 2 and Weibull distribution parameters of Equation 1 via an Expectation-Maximization procedure. A recent study [16] has demonstrated that a vector approximate message passing framework can be applied to imputed and whole-genome sequence (WGS) data to achieve state-of-the-art polygenic risk score performance for human complex traits. Here, we develop this framework further for age-at-onset phenotypes, where the majority of the data is right-censored. The Weibull model we employ, whose applicability to omics and medical data is well documented in the literature [17–19], is highly non-linear and we: *(i)* introduce a novel Weibull channel denoiser, and *(ii)* a novel method for handling right-censored data. These properties make vampW broadly applicable to biological datasets integrated with electronic medical records containing information on disease onset times.

Within a single framework, vampW performs variable selection using Posterior Inclusion Probabilities (PIP, see Methods), while simultaneously enabling time-to-event prediction. Here, we apply this model to analyze protein abundance measurements from the UK Biobank Pharma Proteomics Project (PPP) dataset [3] paired with age-at-onset phenotypes. As compared to the marginal CoxPH testing that has previously been used for these data [2, 4, 5], the key differences are that within vampW: *(i)* all proteins are modeled jointly, capturing combined effects of each plasma protein conditionally on the others; *(ii)* an interpretable hazard structure is used where depending on the value of its shape parameter, the hazard rate can increase, decrease, or remain constant over time; *(iii)* a full-likelihood exploits the actual event times directly, capturing the probability of the observed data under the complete model specification; *(iv)* a continuous survival curve can generate long-term survival predictions because its parametric form is defined over the full time domain, whereas the Cox model paired with the Breslow estimator yields a piecewise-defined baseline hazard; and *(v)* prior knowledge or specific information can be incorporated within the Bayesian framework and posterior mean estimates are produced together with credible intervals, allowing for explicit quantification of uncertainty.

### Simulation study in UKB PPP

We begin by benchmarking vampW against several state-of-the-art variable selection approaches within a semi-synthetic simulation study. Using the 2,924 observed protein levels from the UKB PPP dataset measured in 53,018 individuals, we simulate artificial phenotypic outcomes in 24 scenarios (see Methods for details). The varied parameters are: *(i)* the proportion of variance explained by biological markers, *(ii)* the proportion of censored individuals, *(iii)* the phenotype distribution (Weibull, ExpGamma), *(iv)* the prior distribution of the slab part of the regression coefficients (Normal, Laplace), and *(v)* the ExpGamma distribution parameter. Each scenario is repeated 50 times to obtain independent realizations of the simulated data, and train/test splits are resampled at a 90%/10% ratio across these realizations. We evaluate vampW against several approaches: one-protein-at-a-time testing (CoxPH marginal); non-penalized CoxPH fitted to all markers jointly (CoxPH); joint LASSO-penalized CoxPH with cross-validation and stability selection (CoxPH LASSO); and the deep learning approach DeepSurv (see Methods for details). We assess the ability of each method to recover the simulated causal variables and the out-of-sample time-to-event observation prediction accuracy. For variable selection, we report the false discovery rate (FDR) and true positive rate (TPR) in 4 representative scenarios in Figure 1, where FDR = FP / (FP + TP) and TPR = TP / (TP + FN), with FP denoting number of false discoveries, TP the number of true positive discoveries, and FN the number of false negative discoveries. FDR–TPR plots for all 24 simulation scenarios are reported in Supplementary Figure 1.

**Fig. 1:**
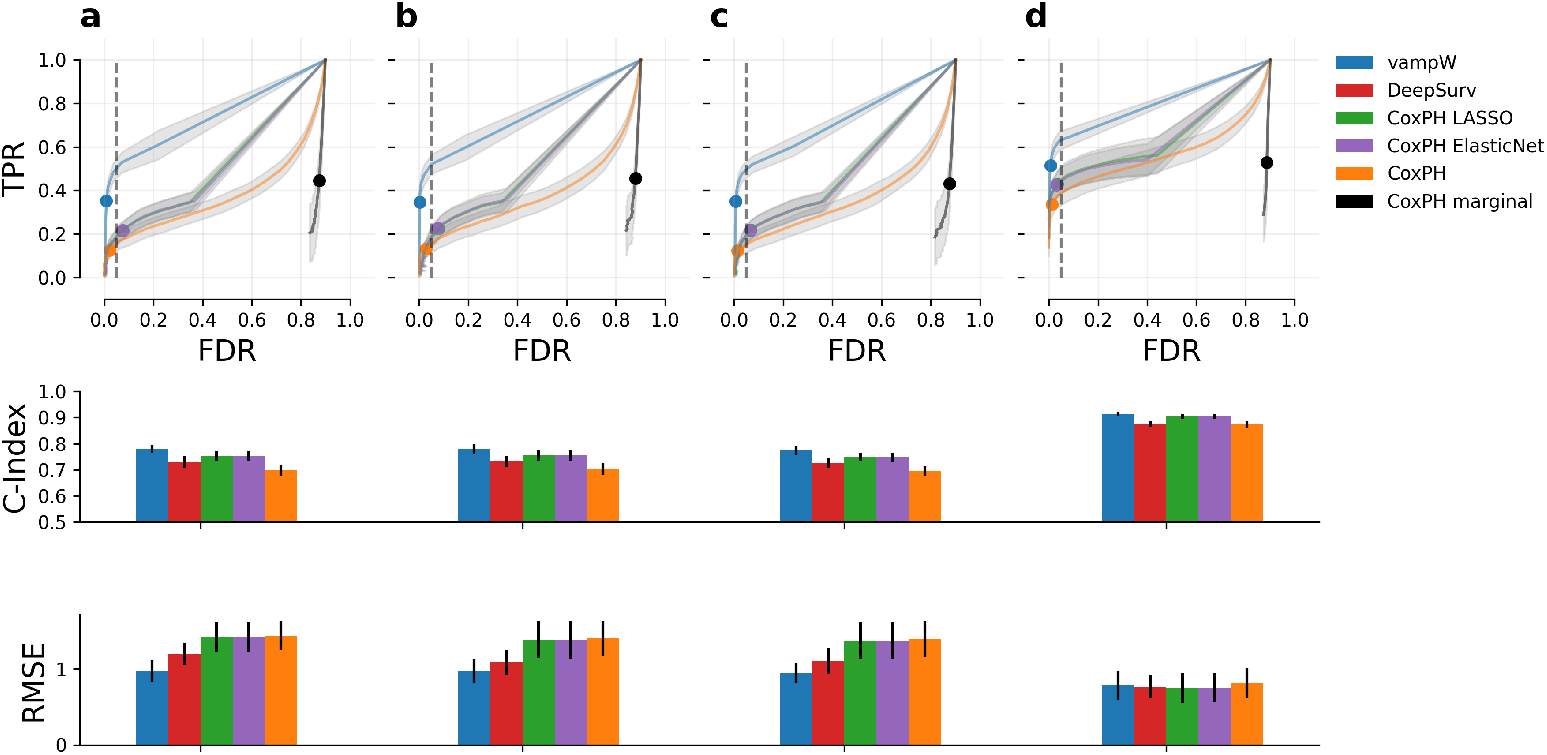
Simulation study of variable selection and out-of-sample prediction performance using 2,924 proteins across 53,018 individuals from the UK Biobank. We present four representative simulation scenarios (out of the 24 in total presented in Supplementary Figures 1-3), covering low signal-to-noise ratios, model mis-specification, and high censoring rates. Specifically, subplots (a, b, c, d) correspond to simulation scenarios (m, o, q, t), as described in Supplementary Table 1. Across all subplots, we use the following color convention: vampW in blue, marginal CoxPH in black, joint CoxPH in orange, LASSO-penalized CoxPH in green, Elastic Net-penalized CoxPH in purple and DeepSurv in red. In the first row, we show variable selection performance as the relationship between false discovery rate (FDR = FP / (FP + TP)) and true positive rate (TPR = TP / (TP + FN)). For vampW, we use posterior inclusion probabilities (PIP), while for the other methods, p-value testing across multiple thresholds is employed, see Methods for details. Colored dots indicate the 0.95 threshold for vampW PIP and the Bonferroni-corrected 0.05 threshold for the other methods. The second row presents prediction performance measured by the concordance index (C-index) on a held-out test set. The last row presents prediction performance measured by the Root Mean Squared Error (RMSE) between the true and predicted survival times in the logarithmic domain, on uncensored individuals. The error bars (black lines in the bar plots and gray shaded areas in FDR-TPR plots) represent the standard deviation across 50 simulation replicates.

The results of Figure 1 show that joint variable selection under the Weibull model of our vampW approach achieves a calibrated false discovery rate, while outperforming both marginal and joint CoxPH models in terms of true positive rate. More precisely, marginal oneat-a-time association testing using the Cox model produces over 80% false positives across all simulated scenarios, even with a Bonferroni correction. This is due to the fact that, when testing each protein individually, correlations can inflate false discoveries as the estimated effect is confounded by other correlated proteins. Modeling the data using a single joint CoxPH model substantially reduces false positives, leading to well-calibrated association testing with a false discovery rate below 0.05%. In contrast, a joint approach reduces the pool of candidate markers for clinical testing while capturing combined or conditional effects, which may be more informative for understanding biological pathways. Furthermore, employing a LASSO-penalized CoxPH model with 5-fold cross-validation and stability selection increases power over the joint CoxPH model, while maintaining a well-calibrated false discovery rate in most scenarios. However, the TPR achieved by the LASSO-penalized CoxPH model with stability selection is still lower than that of vampW across all settings. The complete results for all the 24 simulation scenarios in Supplementary Figure 1 are similar to those presented in Figure 1, where overall, the TPR of vampW is higher, with comparable or improved FDR to CoxPH approaches. This holds even under model mis-specification (generalized Gamma distribution with shape parameter *κ* ∈ {0.8, 1.2}) or when the prior distribution of simulated regression coefficients follows a spike-and-slab prior with slab component being a Laplace distribution which has heavier tails than the Gaussian slab prior used by vampW (Supplementary Figure 1). We investigate the ability of vampW to recover signals from highly correlated proteins, by annotating proteins based on their correlation structure (the top 100 correlated proteins are shown in Supplementary Figure 4). If the correlation of a particular protein with any other protein is greater than 0.5 in terms of the absolute value of the Pearson correlation coefficient, it is categorized as a highly correlated protein, with remaining proteins categorized as low-correlated. Supplementary Figure 5 shows that, among true-positive predictions with PIP ≥ 0.95, the fraction of highly versus low-correlated proteins closely matches the corresponding fraction among ground-truth in the simulation. Thus, vampW has equivalent detection power across proteins of different correlation structure.

We then compare the out-of-sample prediction accuracy across approaches, using two metrics: *(i)* Harrell’s concordance index (C-index) [20], which measures the proportion of concordant pairs, defined as C-index = (*n*_*c*_ · 0.5 *n*_*t*_) / *n*_*a*_, where *n*_*c*_ is the number of pairs which are in the same order in predictions in comparison to the observation, *n*_*t*_ is the number of tied pairs, and *n*_*a*_ is all possible admissible pairs; and *(ii)* root mean squared error (RMSE), which represents the average error between the predicted times and the actual diagnosis times for uncensored individuals, defined as RMSE 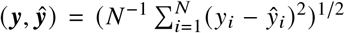, where *ŷ* is the predicted outcome time and (***y***) is the ground-truth for the uncensored individuals. For the simulation results (Figure 1 and Supplementary Figures 2 and 3), the RMSE is computed on log ***y***, which is the quantity standardized to zero mean and unit variance by all methods. For the results on real data (Figure 3), the RMSE is computed on ***y***, which represents the actual time of onset and hence provides a more interpretable quantity.

Prediction accuracies for different methods are presented in Figure 1 for 4 representative scenarios, in Supplementary Figure 2 for all 24 simulation scenarios and in Supplementary Figure 3 considering pairwise differences of these performance metrics across different simulation replicas. These plots show that, in comparison to CoxPH LASSO, vampW achieves higher C-index values across all simulation scenarios, as well as lower or on par RMSE in all scenarios. We additionally compare vampW to DeepSurv, a deep learning extension of the CoxPH model developed for risk prediction. DeepSurv performs better than the standard CoxPH, but is outperformed by vampW in all simulation scenarios in terms of C-index. In terms of RMSE, in most simulation scenarios vampW and DeepSurv exhibit similar performance, with vampW being significantly better in three scenarios. We refrain from comparing association testing performance with DeepSurv, as the latter was primarily developed for prediction tasks (where it does not improve upon vampW) and it does not provide variable selection within the current framework. We note that, in the four simulation settings presented in Figure 1, CoxPH with Elastic Net penalization performs on par with LASSO-penalized CoxPH in both variable selection and prediction. Thus, we do not include it in the additional results of Supplementary Figure 1, Supplementary Figure 2, and Supplementary Figure 3.

To further evaluate the vampW algorithm under more severe model misspecification, we conduct an ablation study that accommodates non-linear effects, interactions, and competing risks. For non-linear effects, we report variable selection and out-of-sample prediction results in Supplementary Figure 6 for four scenarios, corresponding to the simulation parameterizations (*m, o, q, t*) in Supplementary Table 1. In variable selection, vampW outperforms all methods. In the C-index, vampW also outperforms all methods, and falls within DeepSurv’s error bars in three out of four scenarios. In RMSE, vampW and DeepSurv perform comparably, particularly in the high signal-to-noise scenario. We report simulation results with interaction effects in Supplementary Figure 7. In variable selection and C-index, vampW again outperforms the competing methods, while in the high-signal Weibull scenario, vampW and Deepsurv fall within the same RMSE error bars. Finally, Supplementary Figure 8 reports the resulting performance for competing risk simulations. In setups where the simulations follow a Weibull distribution (a,d), vampW outperforms all other methods in terms of RMSE, while in setups where onset times are described with the ExpGamma relationship (b,c), vampW stays competitive in terms of the C-index metric but is outperformed by other methods in terms of RMSE. Thus, in most settings, vampW outperforms or, at least, remains competitive with all other methods in terms of variable selection performance, C-index and RMSE, even when the underlying mechanism accommodates non-linear effects, protein–protein interactions, and competing risks.

### Analyzing 24 disease-related outcomes in UK Biobank

We apply vampW to the age-at-diagnosis of 24 disease outcomes derived from medical records in the UK Biobank, linked to protein measurements from blood samples. In Figure 2, we report two sets of vampW discoveries, passing a PIP threshold of 0.95, for models with and without exponential age correction of the plasma protein levels. First, for the model without exponential age correction, we also run marginal CoxPH, joint CoxPH, and LASSO-penalized CoxPH analyses, all using Bonferroni-corrected p-value testing. Marginal testing identifies tens of times more associations compared with joint vampW testing, consistently across most traits, with exceptions being Alzheimer’s disease, endometriosis, Parkinson disease, vascular dementia, brain/CNS disorders, colorectal and prostate cancer (Supplementary Table 2). However, our simulation study indicates substantially inflated false discoveries when using marginal testing, which often selects features that appear significant individually but are correlated with each other, whereas joint testing produces a potentially more biologically meaningful set of discoveries, reducing the need for unnecessary follow-up experiments. The set of vampW discoveries is largely independent of the top-*x* marginal discoveries, where *x* corresponds to the number of vampW discoveries. Specifically, the overlap between vampW and the top-*x* marginal CoxPH results is 37 out of 219 vampW associations (Supplementary Table 3). This indicates that vampW identifies largely novel associations, and that its joint testing capability cannot be replicated simply by tightening the marginal testing p-value threshold. In comparison with joint CoxPH and LASSO-penalized CoxPH, 61 and 110 associations, respectively, overlap with vampW (Supplementary Table 4). The overlap indicates that vampW is not overly conservative in variable selection, as it identifies proteins that are also detected by LASSO. For 9 of the 24 traits, no association passes the PIP threshold.

**Fig. 2:**
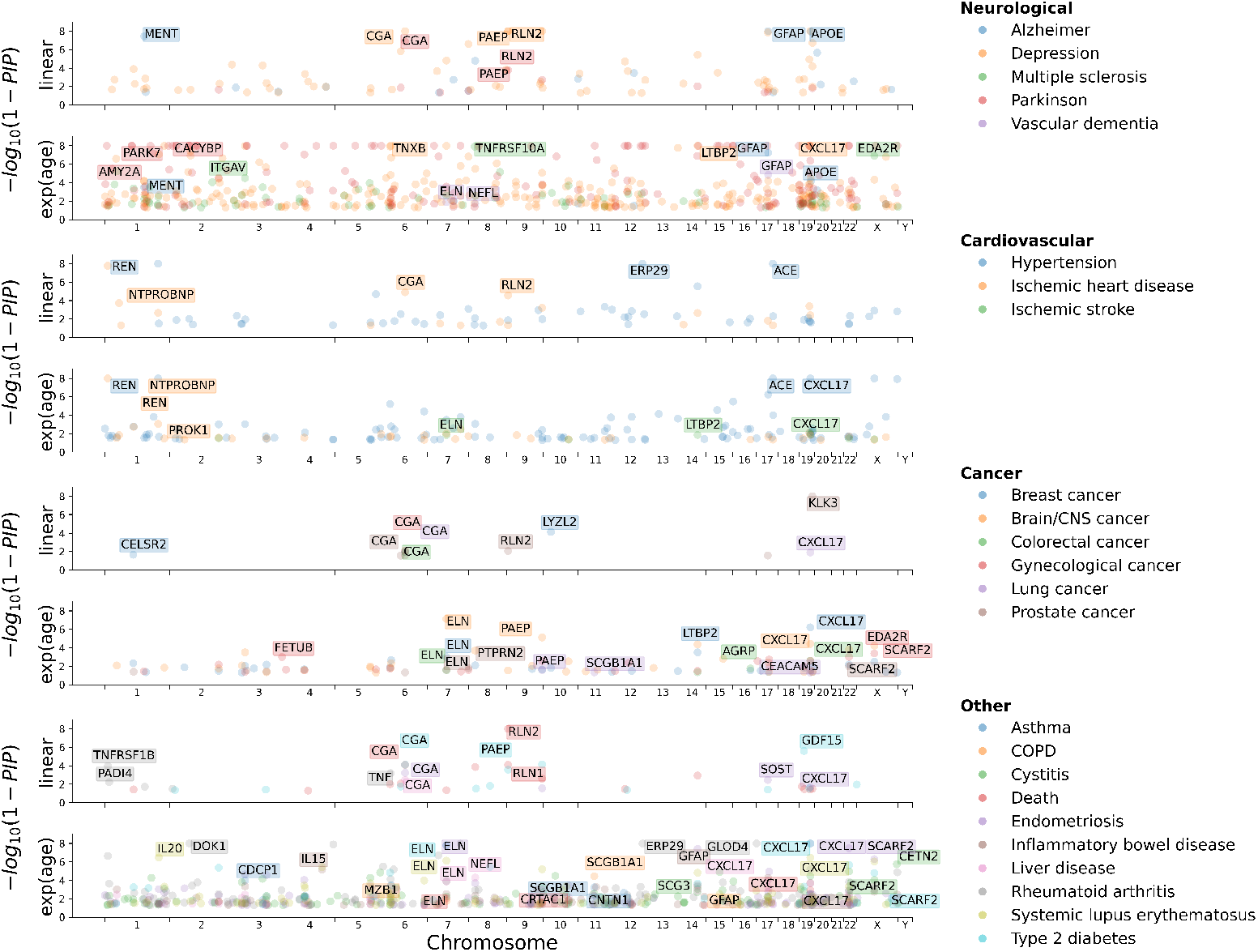
Protein-coding genes discovered by vampW and their genomic positions. The proteins discovered by vampW and their corresponding protein names that pass a posterior inclusion probability (PIP) threshold of 0.95 are shown. Disease outcomes are categorized into four groups: neurological diseases, cardiovascular diseases, cancer types, and other diseaserelated outcomes.

**Fig. 3:**
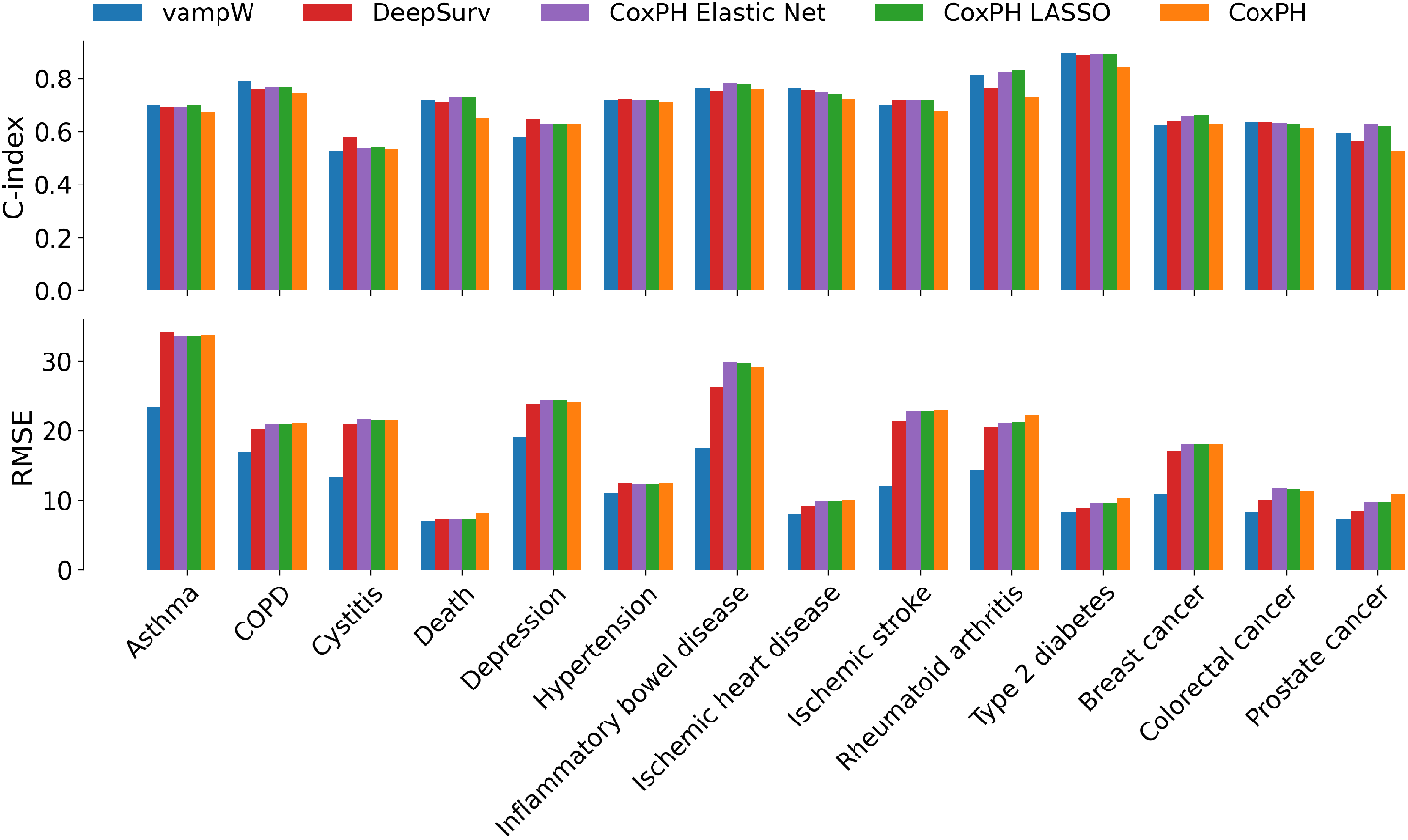
Out-of-sample performance comparison for CoxPH, LASSO-penalized CoxPH, DeepSurv, and vampW. We evaluate time-to-diagnosis predictions for 14 health-related outcomes in the UK Biobank (traits with ≥ 100 cases in the test set) using protein measurements adjusted for age and sex. Model accuracy is assessed using the Concordance Index (C-index), which measures the probability of correctly ranking patient risks, and the root mean squared error (RMSE), representing the average error in years between predicted and actual diagnosis times for uncensored individuals. Training is performed on data that is first log-transformed and standardized to zero mean and unit variance, and then scaled back using an exponential transformation. To represent actual survival times, we rescale the predictions back to years using statistics from the training set. A sensitivity analysis shows that using test-set statistics for this rescaling impacts results negligibly and preserves the ordering of methods in terms of their performance. In terms of RMSE, vampW significantly improves over all other methods. In terms of C-index, vampW outperforms CoxPH, and it performs on par with DeepSurv and CoxPH with LASSO and Elastic Net penalty.

In the set of discoveries without exponential age correction, we highlight the discovery of collagen alpha-1(IX) *COL9A1* in type 2 diabetes, which was also identified by the standard joint CoxPH model but not by CoxPH LASSO. Although it has previously been reported among top plasma proteins [5], it is not well established as a biomarker and may provide novel insights into the biological pathways underlying the disease. Another potentially novel association with type 2 diabetes is the *MATN3* gene, which also exhibits an underlying interaction with *COL9A1* based on text mining, as shown in Supplementary Figure 9. Furthermore, we identify *SOST* as a novel candidate associated with endometriosis, a potential biomarker that has not been widely studied in this context.

Without exponential age correction of the protein data, one of the leading associations we consistently identify is the protein-coding gene *CGA*, which is associated with multiple diseases, including endometriosis, T2D, depression, ischemic heart disease, liver disease, Parkinson’s disease, gynecological cancer, prostate cancer, and colorectal cancer. Notably, plasma levels of *CGA* are highly influenced by age and sex in non-linear ways and rank among the top six protein-coding genes (alongside *FSHB, SOST, GDF15, MLN*, and *PTN*) exhibiting the most significant age-dependent changes [5, 21]. This suggests that plasma levels of these proteins may be associated with these outcomes simply because the proteins and the time-to-diagnosis both vary exponentially with age.

To explore this further, we conduct an additional adjustment of the plasma protein observations including exponential age effects along with the linear ones (see Methods), and re-analyze the traits. This yields associations of plasma protein levels with disease onset beyond age-related variation. The protein-coding gene *CGA*, previously reported as age-associated, disappears from ischemic heart disease, colorectal disease, and mortality associations when non-linear effects of age on protein levels are taken into account (see Figure 2 and Supplementary Figure 10). Moreover, for certain outcomes, the signal becomes attributed to a different set of proteins, revealing new underlying biological structure. For example, we find the chemokine *CXCL17* as associated with colorectal cancer, which has previously been reported as a potential biomarker for the diagnosis of colon cancer [22]. Furthermore, we identify Tenascin-XB (*TNXB*) protein, encoded by the *TNXB* gene, which is recognized as a potential blood-based plasma biomarker associated with depression [23]. *PARK7* (also known as DJ-1) is a multifunctional protein that acts as an antioxidant sensor. Its oxidized form (oxDJ-1) in blood has been proposed as a promising biomarker for early-stage Parkinson’s disease [24]. It was also identified by vampW, after exponential age correction, in association with Parkinson’s disease and rheumatoid arthritis.

Regardless of the age correction approach, we find several associations supported by recent literature. In particular, we find *ACE* to be associated with both hypertension and ischemic heart disease, supporting a role in blood pressure regulation [25]. Prostate-specific antigen (PSA) is the most widely used clinical biomarker for identifying men at risk of prostate cancer prior to symptom onset, and its mechanisms have been extensively studied [26]. PSA is encoded by the *KLK3* gene, which exhibits an association with prostate cancer, regardless of age correction approach in our study. Growth differentiation factor 15 (*GDF15*) reduces food intake and represents a promising therapeutic target for the treatment of type 2 diabetes [27]. It is also a useful biomarker for identifying individuals at increased risk of diabetes [27], and vampW exhibits a strong association between *GDF15* protein levels and type 2 diabetes. Variation in the *APOE* protein, especially the allele *ε*_4_, has been widely established as a risk factor for lateonset disease across multiple large cohorts [28]. Regardless of adjustment for non-linear age effects, we find *APOE* to be significantly associated with Alzheimer’s disease progression. We also identify renin (*REN*) associated with hypertension, which is a key enzyme in the reninangiotensin-aldosterone system (RAAS), making its measurement—often via plasma renin activity (PRA)—a crucial biomarker for classifying, diagnosing, and assessing cardiovascular risk in hypertension patients [29].

We further identify 262 and 9 protein associations, respectively, for the models with and without exponential age correction that show no prior link to a given disease in the DISEASES 2.0 database, which integrates disease–gene associations from automatic text mining and is updated on a weekly basis (we use the March 2026 version). We examine the total singlenucleotide polymorphism (SNP) heritability of proteins, as shown in Supplementary Figure 11, using values reported in [3]. We find no significant deviation of vampW-discovered proteins compared to all proteins in the study, with average SNP heritability of 19.9% and 18.7% for models with and without exponential age correction, respectively. Notably, we identify 25 and 233 proteins, without and with exponential age correction, respectively, that are associated with multiple diseases simultaneously, indicating potential competing risks.

To further validate the vampW discoveries and assess whether they can be replicated in a cohort using a different sampling technology, we matched proteins and outcomes between the UK Biobank and a SomaScan v4 study of plasma samples from 36,000 Icelandic individuals [30]. Among the vampW discoveries, we identified matching proteins for Alzheimer’s disease, depression, hypertension, rheumatoid arthritis, and prostate cancer. Of the 71 matched discoveries, 48 also show an association from logistic regression with binary disease occurrence in the Icelandic cohort at a Bonferroni-adjusted p-value threshold. Moreover, we analyze the protein–disease associations discovered by vampW without exponential age correction in the independent Generation Scotland (GS) study, which used the mass spectrometry technology. We observe limited overlap in the proteins measured across the two studies, and the corresponding p-values for proteins and diseases in the overlap are shown in Supplementary Figure 12. Using a Bonferroni-corrected p-value threshold of 0.1, we replicated 4 out of 13 tested associations, including the association of *CDH5* with hypertension, which is a potentially novel protein biomarker recently recognized as a key contributor to hypertension pathogenesis. [31].

In Figure 3, we show the absolute values of C-index and RMSE metrics across traits with more than 100 uncensored individuals (representing cases) present in the test set. In terms of RMSE, vampW outperforms all other methods across all traits: on average, it reduces the RMSE by 31% with respect to CoxPH, by 30% with respect to both CoxPH LASSO and Elastic Net and by 26% with respect to DeepSurv. Figure 3 gives the RMSE in the actual scale of years.

In terms of out-of-sample C-index performance, vampW shows an improvement over CoxPH in 11 / 14 traits, over LASSO-penalized CoxPH in 6 / 14 traits and over DeepSurv in 9 / 14 traits. Furthermore, on average, the C-index of vampW is 0.5% better than that of DeepSurv, 4% better than that of CoxPH, and 1.3% worse than that of LASSO-penalized CoxPH. Finally, we note that the largest deviations from the Weibull distribution, as shown in Supplementary Figure 13, occur for asthma and inflammatory bowel disease, for which vampW has limited power for association testing but still demonstrates improved time-prediction performance over the various baselines. In comparison to other methods, vampW yields better prediction performance at later ages for most outcomes, based on the Brier score, as shown in Supplementary Figure 14.

We find that vampW generates highly transferable protein risk scores across diverse genetic ancestries, maintaining predictive utility in minority populations. Specifically, while vampW achieves comparable accuracy in terms of C-index to CoxPH LASSO, it consistently outperforms both standard CoxPH and CoxPH LASSO models in minimizing overall prediction error (RMSE), as seen in Supplementary Figure 15. We also conduct an additional ablation, restricting the analysis to incident cases only for two traits with sufficient sample size (hypertension and ischemic heart disease). In Supplementary Figure 16, we show that under this modeling choice, vampW remains competitive in C-index and significantly outperforms the other methods in RMSE. Compared with the original age-at-onset analysis of prevalent and incident cases, the loss of sample size in the incidence analysis causes vampW’s C-index performance to drop by 12.61% for hypertension and 12.97% for ischemic heart disease.

To demonstrate the wider applicability of our algorithm outside of the proteomics domain, we additionally benchmark vampW in data-scarce regimes using common survival benchmarking datasets, following the study [32]. As shown in Supplementary Figure 17, vampW outperforms all deep learning methods on the METABRIC, SUPPORT, and SAC3 datasets, and outperforms 6 out of 13 benchmarking methods on the GBSG dataset.

## Discussion

In this work, we develop the novel vampW algorithm for Bayesian variable selection under the Weibull model. Applying our approach to the largest proteomics data available to date, we demonstrate its capability to jointly fit thousands of plasma proteomics markers across tens of thousands of individuals from the UK Biobank, revealing new disease biomarkers. Our simulation study shows that vampW outperforms state-of-the-art methods for variable selection, reducing false discoveries compared to marginal testing and improving the true positive rate relative to multiple forms of CoxPH models. When analyzing real health-related outcomes from the UK Biobank, vampW identifies 1308 and 219 associations between protein levels and human complex diseases, for models with and without exponential age correction.

The models exhibit a 69% and 68% replication rate in an independent Icelandic cohort using a different technology, accounting for overlap in proteins and outcomes between the studies. The replication of our findings across different technologies is further supported by analysis in Generation Scotland proteomics data.

For out-of-sample prediction of the outcome ranking (C-index), vampW outperforms the standard CoxPH model and achieves performance comparable to penalized versions of CoxPH and to the deep learning-based method DeepSurv. For the prediction of disease onset times in uncensored individuals, vampW yields more accurate predictions, measured by RMSE, as compared to all other benchmarked methods. We also show that vampW generates highly transferable protein risk scores across diverse genetic ancestries and achieves strong out-of-sample prediction performance in other non-proteomics data-scarce regimes, outperforming deep learning methods on most of the examined survival benchmarking datasets, even when trained on very small sample sizes. Moreover, we show that vampW remains competitive with deep learning, even when the underlying mechanism accommodates non-linear effects, protein–protein interactions, and competing risks.

There are several limitations and potential avenues for future work. First, plasma protein levels, as well as other multi-omics markers, may be influenced by multiple confounding factors, such as medication use, sex, and age, including non-linear effects of age on protein levels, as suggested by our ablation study. In this context, a key direction for future work is to leverage the medical records of individuals in the UK Biobank, which would allow adjustments for a broader set of potential confounders. Moreover, to examine whether variation in protein levels reflects disease predisposition or the disease state itself, one possible approach is to restrict the analysis to individuals who experience events after the date of blood sampling and assessment, rather than incorporating both prospective and historical events. With more samples available, covering a larger portion of the population prevalent with a certain disease, studying associations between survival time and protein levels during the follow-up period represents a viable approach. In our ablation study, using two medical outcomes with sufficient numbers of follow-up incident cases, we demonstrated the utility of the vampW framework for this type of modeling choice. Additional extensions are models that can identify relevant agerelated changes in protein levels that are associated with disease onset at different time points, which requires repeated proteomics observations. Another avenue for future improvement would be to model multiple disease outcomes jointly while correcting for competing risks, allowing the identification of disease-specific risk rather than broader biological vulnerability, as well as to improve RMSE performance for competing risks themselves via a different likelihood modeling. Finally, our future work will also focus on improving the computational efficiency of the vampW algorithm through parallelization and reduced memory requirements, which has the potential to enable large-scale analyses involving millions of genomic markers.

In summary, vampW is a novel approach for selecting multiomic biomarkers associated with human diseases and for predicting disease onset times. With increasing sample sizes and expanding data coverage in the world’s largest biobanks, joint vampW analyses have the potential to advance our understanding of the genomic architecture of complex human diseases by enabling interpretable identification of disease associations and providing accurate predictions of disease onset.

## Methods

### Weibull model for time-to-event outcomes

We consider a Weibull (*α, η*) model, one of the most common models in survival analysis, where *α* and *η* are the shape and scale parameters. We follow a recently proposed formulation of the model [17], where we aim to estimate mean and variance of the time-to-event outcome. Let *y*_*i*_ be the time at which the individual *i* experiences the event, and consider the survival function 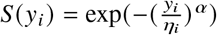. In the logarithmic space, the mean and variance of the model can be separated as follows:

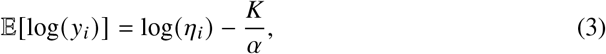

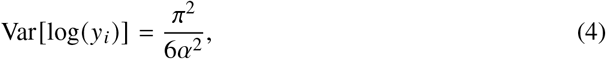

where *K* is the Euler-Mascheroni constant (≈ 0.57722). Once the variance is independent of the scale parameter *η*_*i*_, one can introduce the vector of underlying latent genetic effects ***β*** as 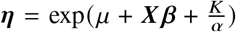, resulting in E log ***y*** = *μ* ***X*** + ***β***, where ***X*** ∈ ℝ^*N* ×*P*^ is a design matrix containing observations on *N* individuals and *P* biological markers, ***y*** = [*y*_1_, …, *y*_*N*_], ***η*** = [*η*_1_, …, *η*_*N*_], ***β*** is the vector of marker effects, and *μ* represents the model intercept. The columns of the matrix ***X*** are normalized, i.e., they each have zero mean and unit variance. Using the fact that the logarithm of a Weibull random variable follows a Gumbel distribution and the Location-Scale property of the Gumbel distribution, we formulate the final model as

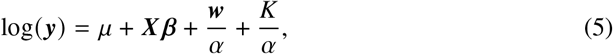

where ***⊤*** = [*w*_1_, …, *w*_*N*_] with each *w*_*i*_ following the Gumbel (0, 1) distribution. We note that in the algorithm and software described below, any arbitrary measurement represented in floating point can be accommodated as the entries of ***X***. These can be gene expression, methylation, copy number variants, DNA sequence variation, protein measures, which can all be included alongside one another.

To allow for a range of effect sizes from different data types, we select the prior on the marker effects ***β*** to be distributed as a form of spike and slab distribution, with non-zero signals coming from a mixture of normal distributions,

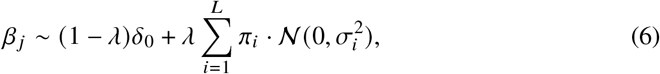

where (*π*_1_, …, *π*_*L*_) denotes the vector of inclusion probabilities for each of the *L* mixture components, *δ*_0_ is a Dirac spike function at 0, *λ* is the sparsity ratio, and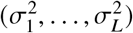 are mixture-specific variances.

Finally, we account for the right-censoring issue inherent in time-to-event data. In practical applications, the true event time *y*_*i*_ is often right-censored, meaning that the event of interest has not occurred for some individuals by the end of the study. Consequently, we do not observe *y*_*i*_ directly for all *i*; instead, we observe the time min (*y*_*i*_, *C*_*i*_) and an indicator *δ*_*i*_ = I (*y*_*i*_ ≤ *C*_*i*_), where *C*_*i*_ is the censoring time and *δ*_*i*_ = 1 implies the event was observed while *δ*_*i*_ = 0 implies it was censored.

### vampW algorithm and its implementation

To infer the parameters of Equations 5 and 6, we propose an algorithm that we call vampW, which is based on Vector Approximate Message Passing (VAMP) for generalized linear models [11, 12], where the outcome variable is observed through a noisy and non-linear Weibull channel. It adaptively estimates the prior-specific parameters 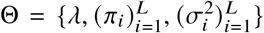 via a Bayesian Expectation-Maximization procedure, as introduced in EM-VAMP [33]. vampW introduces a novel approach for handling censored data, as well as denoisers derived specifically for Weibull link functions and the corresponding expectation-maximization updates. We highlight that no existing AMP method is able to address the challenge of censored data.

As seen in Algorithm 1, vampW consists of two denoising steps and two (non-linear) minimum mean square error (MMSE) estimation steps, applied separately to the vector of effect sizes ***β*** and to the latent predictor. The denoising step accounts for the prior properties of the model when considering a noisy estimate of either ***β*** or the latent predictor, while the MMSE step further refines the estimates by taking the correlation structure among the columns in the design matrix ***X*** into account. The key feature is the so-called *Onsager correction*, which is added to ensure the asymptotic normality of the noise corrupting the algorithm’s estimates of ***β*** at every iteration, important for the subsequent feature selection. Following [16], we employ the Hutchinson estimator [34] to evaluate the trace of the inverse of regularized Gram matrices, and we warm-start the conjugate gradient (CG) algorithm approximating the solution of the linear system (line 32 in Algorithm 1). The CG solver is initialized from the value at convergence in the previous iteration, which leads to reduced computational time. The CG solver providing the initial guess for the non-linear LMMSE equation is warm-started from the previous iteration’s 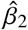, and the equation is then solved with `scipy.optimize.fsolv` using the analytic Jacobian. In the Hutchinson estimator, all eight probes vectors are solved jointly via a single Cholesky factorization of the Jacobian. Compared to the gVAMP algorithm introduced by [16] for a linear model without censoring, vampW derives a non-linear Weibull model within the AMP framework and incorporates censoring information into the process (as detailed in the paragraph below), thus requiring two denoising steps and two non-linear MMSE estimation steps.

A direct application of generalized EM-VAMP to real genomics, epigenetics and proteomics measures may lead to signal-estimation divergence, primarily due to violations of right-rotational invariance in the data design matrix, one of the assumptions in the original EM-VAMP algorithm. To mitigate these stability issues, we perform *damping* of denoised signals, as seen on line 11 of Algorithm 1, where *ρ* ∈ (0, 1) denotes the damping factor. This approach makes the algorithm take smaller steps when learning ***β***. In order to further stabilize convergence and improve prediction performance, we employ damping autotuning [35]: we iteratively reduce the damping by the factor *c*_*ρ*_ while the training prediction, using signal estimates from the denoising step 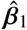, increases over the last iteration, measured by the concordance index (C-index), as seen in line 12 of Algorithm 1. Thus, in terms of model stability, although the data applications described below have design matrices that violate the assumptions required for Bayes optimality of our algorithm, introducing damping and damping autotuning empirically results in strong predictive performance. The damping choices affect the convergence properties when searching over the parameter space, as too strong damping may cause the algorithm to get stuck in local minima; and thus we propose a grid search of starting parameters which is facilitated by the efficient implementation of our algorithm.

#### Algorithm 1

vampW

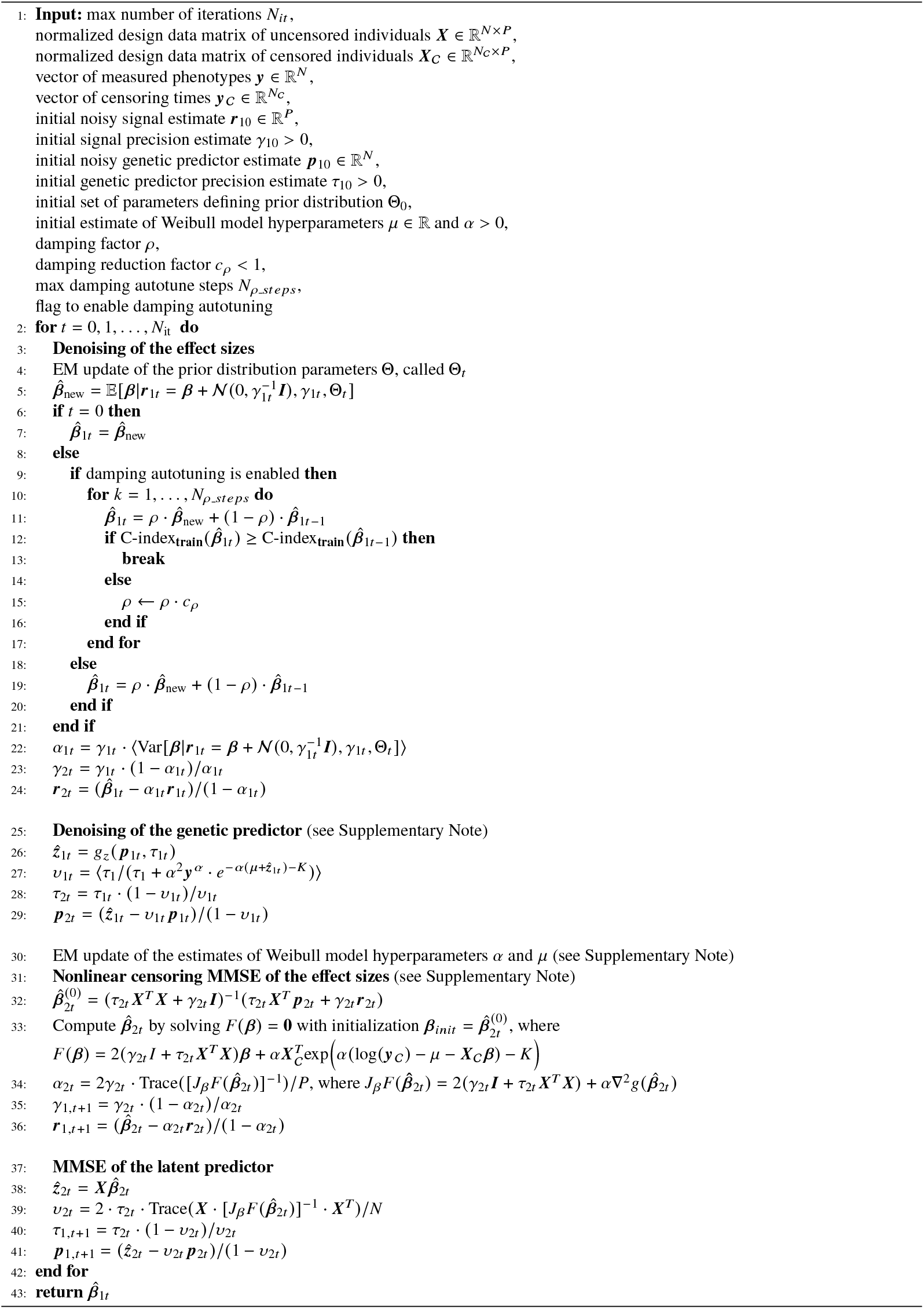

### Handling right censoring with vampW

As disease outcomes are only partially observed, with data being frequently right-censored, we revise the LMMSE formulation of the standard VAMP algorithm for generalized linear models to explicitly incorporate the censored phenotypes. Let = {*i* | *δ*_*i*_ = 0} denote the set of indices corresponding to the *N*_C_ censored individuals 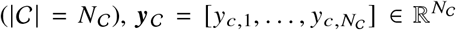 the censored observations and ***z*** = ***X β*** the corresponding linear predictors, where ***X***_*C*_ represents the matrix containing only information on individuals in *C*. We note that both the data matrix and the phenotype vector are split into censored and uncensored individuals at runtime, without the need to perform the splits manually beforehand.

We reformulate the problem by incorporating the survival function of the standard Gumbel distribution *S*_*w*_ into the likelihood term of the optimization problem. We recall that the survival function of the standard Gumbel distribution evaluated at *α* (log (*y*_*c,i*_) − (*μ* ***X***_*C*_ ***β***_*i*_) − *K* reflects the probability that the censored individual *i* survived longer than the censoring time *y*_*c,i*_. Relying on the state evolution result that noisy version of the signal and genetic predictor behave as 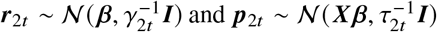, and motivated by the previous interpretation, the final optimization task at hand becomes:

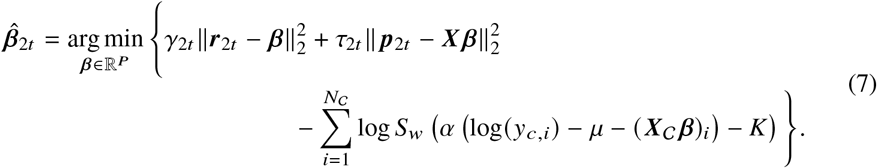

This objective function represents a Maximum A Posteriori (MAP) estimator for the current VAMP iteration. The quadratic terms incorporate the extrinsic information passed from the VAMP factor graph (approximated as Gaussian priors with precisions *γ*_2*t*_ and *τ*_2*t*_), while the logarithmic term explicitly handles the non-Gaussian nature of the censored data by maximizing the survival probability *S* for the unobserved event times. Following the derivation described in the Supplementary Note, we obtain the following formulae for the Onsager correction terms at iteration *t*:

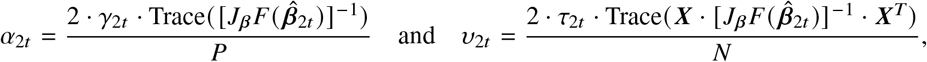

where *F* : ℝ^*P*^ → ℝ^*P*^ is the gradient of the objective function defined as

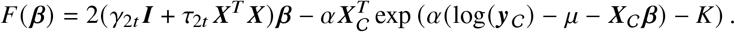

### Association testing

To evaluate the statistical relevance of the markers and test their associations with the traits under investigation, vampW provides Bayesian Posterior Inclusion Probabilities (PIP). Following Algorithm 1, PIPs are calculated using the noisy signal estimates ***r***_1,*t*_ and the associated precision *γ*_1,*t*_ at iteration *t*, relying on the interpretation of those quantities within AMP frameworks in the standard setting with no censoring. We employ the PIP formulation in [16], i.e.,

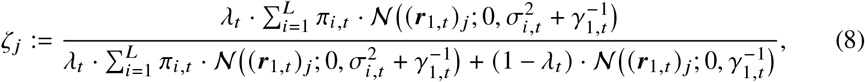

where *ζ*_*j*_ is the posterior probability that marker *j* has a non-zero effect, and 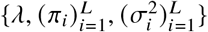 are prior parameters (Equation 6) adaptively learned throughout the vampW iterations. The PIP can be viewed as the Bayesian analogue of a p-value. We use a PIP threshold of 0.95, which represents strong evidence that a variable is associated with the outcome and corresponds to a 5% probability of false discovery in a calibrated setting.

### Computational efficiency

vampW can be run as a single-threaded program on an HPC cluster, requiring approximately 64 GB of random-access memory (RAM) and an average of 4 CPU hours (measured on AMD EPYC 9654 Processor). This corresponds to approximately 0.29 GBP when run on a cloud-computing DNAnexus virtual machine (mem3_ssd3_x8)^1^, which is required due to data protection. We note that in most disease analyses, our stopping criterion (best training C-index) was met around iteration 10, without the need to run the full 30 iterations. The CoxPH LASSO approach can be parallelized, requiring a total of 160 GB of RAM when using 10 parallel CPU cores. With an average of 60.8 CPU hours required per trait, this corresponds to 10.5 GBP per trait when run on a mem3 ssd3 x24 machine. We note that the LASSO runtime depends on the range of penalty parameters explored during the validation step and could therefore be reduced by narrowing this range for specific outcomes. In comparison, CoxPH requires an average of 0.3 CPU hours per trait, and marginal CoxPH requires 0.1 CPU hours per trait, however those methods score lowest in our benchmarks in out-of-sample prediction and variable selection as we show in the Results.

### UK Biobank protein data and time-to-event phenotypes

To validate our proposed approach, we analyze protein measures from the UK Biobank Pharma Proteomics Project (PPP) dataset [3]. We use a total of 2,924 unique proteins measured across 53,018 individuals, obtained using the antibody-based Olink Explore 3072 PEA. We standardize each protein vector across individuals (ignoring missing values) to obtain zero mean and unit variance, and then impute the missing values with zeros. We adjust the protein levels by regressing them on the covariates sex and age via the linear model *α*_1_ · sex + *α*_2_ · age. We then use the resulting residuals, representing the component of protein variation not explained by sex and age. This follows previous studies which use the same correction factors [2, 5], having shown that adjusting for genetic principal components, protein batch, and study center has minimal effects on protein levels. Additionally, to explore non-linear effects of age on protein levels [21], which have not been previously considered in protein-disease onset association testing, we perform a covariate adjustment by including an exponential age component, alongside the linear term, in the model *α*_1_ · sex + *α*_2_ · age + *α*_3_ · exp (age). Throughout all analyses, we use a 90%/10% split for train and test subsets.

The 24 phenotypes selected include: breast cancer, prostate cancer, colorectal cancer, lung cancer, gynecological cancer, brain/CNS cancer, multiple sclerosis, major depression, systemic lupus erythematosus, endometriosis, vascular dementia, amyotrophic lateral sclerosis, inflammatory bowel disease, Alzheimer’s dementia, cystitis, rheumatoid arthritis, Parkinson’s disease, ischemic stroke, COPD, type 2 diabetes, ischemic heart disease, mortality and hypertension, as described in Supplementary Table 5. We choose not to include schizophrenia because preliminary analyses of its age-at-onset distribution reveal a substantial deviation from a Weibull model, along with a comparatively uniform hazard rate across the observed time span, see Supplementary Figure 13. We consider an individual’s time of event from the UK Biobank health records of the first reported occurrence (Category 1712). For major depression, we combine two data fields into one joint set, particularly the first reported depressive episode (UK Biobank code 130894) and the first reported recurrent depressive disorder (UK Biobank code 130896). Similarly, for inflammatory bowel disease, we combine the first reported cases of Crohn’s disease (UK Biobank code 131626) and ulcerative colitis (UK Biobank code 131628) into one joint set. In the type 2 diabetes cohort, we exclude individuals who are observed to have type 1 diabetes (UK Biobank code 130706) at the same time.

For cancer phenotypes (Breast, Prostate, Colorectal, Lung, Gynecological, and Brain/CNS), we use data from the UK Biobank cancer registry. Cases are defined by the presence of disease-specific ICD-10 (field 40006) or ICD-9 (field 40013) codes across any of the 22 reported instances. The specific codes for each cancer type are detailed in Supplementary Table 6. For identified cases, the time-to-event is defined as the minimum age at diagnosis recorded across all matching instances (field 40008). For individuals without a diagnosis (censored observations), the censoring time is calculated as their age at the initial assessment center visit, derived from the date of assessment (field 53) and year of birth (field 34). The age information is derived from calendar years. In the ablation analysis of incident cases, we set the follow-up period to 2023 or the date of death, and model time to disease onset in years.

To assess how population homogeneity affects vampW performance, we repeat our analysis by also partitioning individuals based on their genetic ancestry. Using a Random Forest classifier trained on the 1000 Genomes Project reference panel, we project target samples onto the top six principal components of genetic variation, applying a strict assignment probability threshold of 0.9. From an initial cohort of 53,018 individuals, high-confidence classifications yield 46,011 British or Irish, 1,253 African, and 966 South Asian individuals. A further group of 772 individuals are classified as having other European origin. Quality control and processing of all single nucleotide polymorphisms follow the established pipeline from [36]. Next, we train vampW using only data from individuals of British or Irish descent. To test the model’s performance, we evaluate it against three groups: a British/Irish test set, and the complete sets of African and South Asian participants. Performance is assessed using 10-fold random subsampling, where the evaluation set is repeatedly split into subsets comprising 50% of the original data. For each iteration, we calculate the C-index and RMSE to measure predictive accuracy. vampW operates on the outcome vector ***y*** in the time domain, after standardization on the logarithmic scale, so that log (***y***) has zero mean and unit variance. To obtain root mean square error (RMSE) values in units of years for predictions of UK Biobank outcomes, we rescale the model predictions from the log-standardized space back to the original time scale, using the mean and standard deviation of log (***y***_*train*_), with ***y***_*train*_ denoting the training part of ***y***.

### Replication in Generation Scotland proteomics

#### Generation Scotland cohort

We sought replication of a subset of the vampW findings in an independent dataset, Generation Scotland (GS). GS is an epidemiological study consisting of approximately 24,000 volunteers from across Scotland, aged 17–99 years old at recruitment between 2006 and 2011 [37]. Blood samples were collected via venepuncture during the baseline clinic appointment for over 20,000 volunteers, alongside health, cognitive, and lifestyle questionnaires. Participants consented to electronic-health records linkage to both primary and secondary care data, allowing analyses of incident disease.

#### Mass spectrometry proteomics

Circulating proteins in GS were measured from serum using a high flow-rate liquid chromatography tandem mass spectrometry, which generated data for 325 unique, high abundance proteins across 18,826 participants. The sample preparation process has been described before [38–41]. 5 µL of serum samples were added to a solution of 50 µL of 8 M urea and 0.1 M ammonium bicarbonate at pH 8.0 to have the proteins denatured. Subsequently, the proteins were reduced using 5 µL of 50 mM dithiothreitol for 1 hour at 30 °C and alkylated with 5 µL of 100 mM iodoacetamide for 30 minutes in the dark. The sample was then diluted with 340 µL of 0.1 M ammonium bicarbonate to a concentration of 1.5 M urea. The next step was the trypsinisation of the proteins and 200 µL of the solution was used, and the proteins were digested overnight with trypsin (12.5 µL, 0.1 µg/µL) at 37 °C at a 1/40 trypsin/total protein ratio. The digestion was quenched with the addition of 25 µL of 0.1% v/v formic acid (FA). The peptides were cleaned up with C18 96-well plates, eluted with 50% v/v acetonitrile (ACN), dried by a vacuum concentrator and redissolved in 50 µL of 0.1% v/v formic acid (FA) for processing by liquid chromatography (LC)-MS. Raw data were analyzed by DIA-NN [42] as described previously [41]. For 224 proteins with <20% missingness (i.e., <20% of participants lacked a measured protein level), the missing values were imputed after processing using softImpute method (2.89% imputed values in total). The remaining 101 proteins were subset to individuals with measured protein levels. All proteins were subsequently rank-based inverse normalised prior to downstream analysis.

#### Disease diagnosis information and statistical analysis

Diagnosis information for diseases of interest (Alzheimer’s disease, depression, hypertension, and prostate cancer) was obtained through linkage to primary and secondary healthcare records. If an individual was not diagnosed with a disease, the censor date was set to the most recent date of data linkage (October 2023) or death, resulting in a maximum follow-up period of 17.8 years from baseline appointment. Individuals with a prevalent diagnosis were excluded from the analysis, and the earliest diagnosis was considered. Secondary care records were available for all volunteers. Despite having volunteer consent to access primary care data, these were only accessible for 30% (n = 7,092) of the cohort due to consent constraints with individual GP surgeries (the data holders). All statistical analyses in GS were performed in R v4.4.2 (2024-10-31). Marginal associations between proteins and time-to-disease onset were tested using mixed effects Cox proportional hazard models from the coxme R package (v2.2-22)[43] adjusted for age, sex, and relatedness. Relatedness, modelled with a kinship matrix constructed using the kinship2 R package (v1.9.6.1)[44], was included as a random effect to account for family structure. Only 12 proteins overlapping with vampW findings were analysed. Sample sizes varied across diseases and proteins tested. Alzheimer’s disease: ncases = 105 and ncontrols = 18,709; depression: ncases = 521 and ncontrols = 17,303 except for unimputed B2M protein with >20% missingness (ncases = 352 and ncontrols = 11,064); hypertension: ncases = 1,732 and ncontrols = 15,756 except for unimputed CDH5 protein with >20% missingness (ncases = 1,075 and ncontrols = 9,668); prostate cancer: ncases = 276 and ncontrols = 7,577 as the dataset was restricted to males only. Statistical significance of each protein–disease association was assessed using Wald tests.

#### Simulation study

We conduct a simulation study to demonstrate the utility of the vampW method as a prediction and association-testing tool. We exploit pre-processed (standardized, not adjusted for covariates) real protein measures from the UK Biobank PPP dataset, described above. We simulate artificial phenotypic outcomes in different scenarios, varying multiple parameters, namely: *(i)* the proportion of variance explained by biological markers 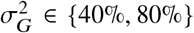, *(ii)* the proportion of censored individuals *C* ∈ {90%, 95%}, *(iii)* the phenotype distribution (Weibull, ExpGamma), *(iv)* the prior distribution of the slab part of the regression coefficients (Normal, Laplace), and *(v)* the ExpGamma distribution parameter *κ* ∈ {0.8, 1.2}. This yields 24 scenarios in total, summarized in Supplementary Table 1. Details on simulating ExpGamma outcomes are in the Supplementary Note.

The causal signals are sampled from either an i.i.d. zero-mean Bernoulli-Gaussian or Bernoulli-Laplace spike and slab distribution:

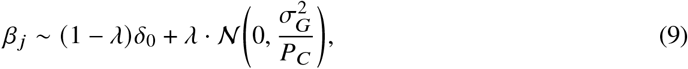

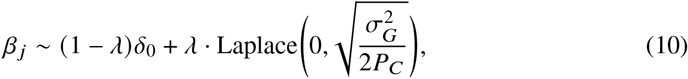

where 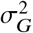 is the target variance explained by biological markers (plasma protein levels). The aim is to spread the variance explained to a certain number of causal markers *P*_*C*_. We use *λ* = 0.1 which, given the total of 2,924 markers, corresponds to roughly 292 causal markers. Note that the variance of the Laplace distribution is determined by its scale parameter. We then generate the phenotypic outcome *y*_*i*_ as either a Weibull or an ExpGamma distribution, as described in the Supplementary Note. In order to extend our simulations to more severe model misclassification, we simulate non-linear effects, interactions and competing risks. For nonlinear effects, we accommodate an underlying signal generation mechanism that combines linear and quadratic term, assigning 50% of the signal to linear effects and 50% to quadratic effects, while retaining the same causal positions. For numerical stability, we clip the extreme values of the quadratic term below the 0.1% and above the 99.9% percentiles. We also simulate interaction effects, in addition to the linear term, assigning 20% of the overall signal to pairwise protein–protein interactions, represented as a multiplication term of two interacting proteins. We select protein interactions from the STRING database [45] with scores above 500. Further, to assess the robustness of our approach in the presence of competing risks, we generate two event times, one for the primary event and one for the censoring process. Both event times are simulated under the same distributional setup, using either a Weibull or an ExpGamma model, with genetic effects drawn from either a Gaussian-slab or a Laplace-slab spike-and-slab prior. At each causal variant, the effects on the two event types are drawn jointly from a bivariate distribution with genetic correlation *ρ* = 0.8, allowing the competing event to share part of the genetic architecture of the primary event. Distribution-specific noise levels are calibrated to achieve the prespecified heritability *h*^2^ for each latent event time. The observed time is defined as *Y* = min (*T*_1_, *T*_2_), with the event indicator set to one when the primary event occurs first and zero when the competing event occurs first.

To simulate data that more closely mimics real-world cohorts, we also account for censoring in the simulation procedure. Specifically, we randomly select a proportion *C* of individuals to be censored. For each censored individual *i*, we multiply the simulated outcome *y*_*i*_ with a random factor drawn from a uniform distribution Unif 0, 1. For the simulation of competing risks, we shift the competing-event time distribution using the empirical quantile of log (*T*_1_) − log (*T*_2_) to attain the target competing-event proportion *C* and standardize the observed log-times using the mean and standard deviation among primary-event observations. We repeat the simulation 50 times for each scenario to obtain independent realizations of the simulated data. We also resample train/test sub-splits in ratio 90%/10% across data realizations.

#### Initialization of vampW

The vampW framework allows several parameters to be specified before running the algorithm. Some of these are estimated from the data, while others influence convergence behavior. In our simulation study, we intentionally initialize vampW with a mis-specified prior to demonstrate that the algorithm can adapt and learn the prior from the data. Following the prior definition in Equation 6, we initialize the slab part of the prior with unit variance and sparsity *λ* = 0.3. While the majority of traits utilize a unit slab part variance, a smaller variance of 0.01 is applied to specific diverged scenarios. In our analysis, we observe better stability of the algorithm when using only 1 slab component (*L* = 1). Furthermore, we initialize the *α* parameter in the Weibull model to 3 (Equation 5). We also postpone learning the *α* parameter until iteration 5 to improve stability. We fix the damping factor at 0.3 and train the algorithm for 30 iterations. Based on our observations, this number of iterations is sufficient for the hyperparameters to stabilize. For simulations with competing risks, we initialize the *α* parameter to 2 and postpone its learning to iteration 4. Finally, we choose the best iteration based on train concordance index after 5 warm up iterations.

We initialize vampW for all 24 UK Biobank traits with a consistent prior structure, setting the initial shape parameter *α* = 3 and sparsity rate *λ* = 0.3 and only one mixture component in the slab part. While the majority of traits utilize a slab variance of 0.01, a wider variance of 1 is applied to specific phenotypes such as hypertension and death. The global shrinkage parameter *ρ* is initialized between 0.3 and 1.0 depending on the trait, with an adaptive *ρ* strategy enabled for a subset of runs, including depression, Parkinson’s disease, and several cancer endpoints, to allow for dynamic adjustment of update speed. More details can be found in Supplementary Tables 7-8. As in the simulations, we choose the final iteration based on the best training concordance index after 5 warm-up iterations. For the model with exponential correction, we initialize vampW with the same priors as in the main analysis, with the exception of depression (initialization label f) and breast cancer (initialization label b), as described in Supplementary Table 7.

### Benchmarking methods

We evaluate vampW against state-of-the-art survival analysis approaches, specifically: *(i)* oneat-a-time marginal CoxPH testing (CoxPH marginal); *(ii)* non-penalized CoxPH fitted to all markers simultaneously (CoxPH); *(iii)* LASSO-penalized CoxPH [8] with cross-validation and stability selection (CoxPH LASSO); and *(iv)* DeepSurv [15]. One-protein-at-a-time testing (CoxPH marginal) is among the most commonly used approaches for variable selection. Joint models offer an alternative which fit each marker in the structural context of all other markers. Furthermore, a penalized model (CoxPH LASSO) enables joint selection of markers while shrinking or zeroing out some coefficients. Cross-validation is used to determine the optimal level of penalization, and stability selection is then applied to identify the most robust predictors and reduce the impact of stochastic variability. DeepSurv represents one of the most commonly used deep learning approaches for survival analysis, capable of modeling complex nonlinear relationships among biological markers. For CoxPH and CoxPH marginal, we use the Python *lifelines* [46] library with its default settings, which estimates the baseline hazard using the Breslow estimator and outputs the regression coefficients and corresponding pvalues from two-sided Wald test. Bonferroni correction is applied for multiple testing rejecting the null hypothesis *H*_0_ if the p-value *p* < 0.05 / *P*, where *P* = 2,924 tests correspond to the number of proteins.

For traits where CoxPH with default parameters failed to converge, we follow the guidelines provided by the *lifelines* documentation for resolving convergence issues in the Cox Proportional Hazard model^2^. Consistent with these recommendations, we use fixed initialization parameters with adjusted penalization and step sizes to ensure stability. Specifically, for liver disease, we employ a penalization of 0.1 and a step size of 0.95; for Alzheimer’s disease, multiple sclerosis, Parkinson’s disease, systemic lupus erythematosus, vascular dementia, as well as the colorectal, gynecological, and lung cancer datasets, we employ a penalization of 0.001 with a step size of 0.95; finally, for inflammatory bowel disease, and brain and CNS cancer, we employ a penalization of 0.001 and a step size of 0.5.

For CoxPH LASSO, we use Python’s *scikit-survival* [47] implementation with a grid search to identify the optimal penalty. We perform 5-fold cross-validation, further splitting the training data, and sweep over 16 penalizer values in logarithmic space, from 10^−4^ to 10^0^. After selecting the optimal penalizer, we perform stability selection [10] by fitting 100 parallel penalized regressions, each trained on a random 75% subset of the data. We then retain only the stable variables appearing in at least 90% of the models. Finally, we fit a non-penalized CoxPH model from *lifelines* using only these stable features, estimating the baseline hazard via the Breslow method to obtain the final refined regression coefficients and their corresponding p-values. Similarly, we train a Cox proportional hazards model with Elastic Net (EN) regularization and stability selection using *CoxnetSurvivalAnalysis* from the *scikit-survival* [47] package. We first perform a grid search over the following L1 penalization ratios, {0.7, 0.9, 0.95, 0.99, 1.0}, and same penalizer values as for CoxPH LASSO, using the package’s default method for estimating the penalization parameter, and select the value based on the best validation performance. Further, we perform stability selection in the same way as described above for the CoxPH LASSO approach.

For running DeepSurv, we use the code provided by the authors [15], using one hidden layer and turning the batch normalization on, and standardization off. To optimize performance, we explore the hyperparameter space via a grid search over the following parameters: ℓ_2_Reg ∈ {0.01, 1, 5}, dropout ∈ {0, 0.1}, learning rate (LR) ∈ {0.001, 0.0001}, size of the hidden layer ∈ {256, 512, 1024}. For non-linear effects, interactions and competing risk simulations, we extend the number of hidden layers with the sizes corresponding to {1024, 512, 256} and ∈ {2048, 1024, 512}, and consider L2 regularization strengths of ∈ {0.01, 0.1, 1, 5}. Moreover, we fix the LR decay to 0.001, momentum for the Adam optimizer to 0.9 and number of epochs to 500. During the grid search, the training set is further randomly split into an 90% sub-train set and a 10% validation subset, optimizing for the best C-index on the validation set. To predict age at onset using DeepSurv, we use the predicted log-risk scores from DeepSurv as input to the *lifelines* CoxPHFitter, which estimates the baseline hazard using the Breslow estimator and predicts expected onset times. For CoxPHFitter, we set the step size to 0.5.

The RMSE is evaluated using only uncensored samples, as the true event times for censored data are unobserved and the benchmarked methods do not explicitly model the censoring time distribution.

#### Benchmarking in data-scarce regime

We conduct a comparison of vampW with deep learning survival methods on several commonly used datasets, namely METABRIC, GBSG, SUPPORT, and SAC3, obtained from the *pycox python* library [48]. Following the study [32], we examine vampW in the data-scarce regime by subsampling the datasets to 100 samples for training and 25 samples for testing. We recreate the exact subsets used in [32], enabling a direct comparison of our results with those reported in the central analysis of the study. We report results for several recent and more traditional deep learning methods, namely: Multi-Task Logistic Regression (MTLR) [49], DeepHit [50], DeepSurv [15], LogisticHazard [51], CoxTime [52], CoxCC [52], PMF [53], PCHazard [53], BCESurv [54], DySurv [55], SumoNet [56], DQS [57], and NeuralSurv [32]. Here, we initialize the vampW prior with one slab component of unit variance and 50% probability. We set the initial parameter *α* to 3 and the damping factor to 0.3, allowing for automatic damping tuning over a maximum of 4 iterations.

## Acknowledgments

We thank members of the Medical Genomics group at ISTA for their comments, which improved this manuscript. We would like to acknowledge the participants and investigators of the UK Biobank study. Generation Scotland was funded by a grant from the Chief Scientist Office of the Scottish Government Health Directorates (CZD/16/6) and the Scottish Funding Council (HR03006). Genotyping and DNA methylation profiling of the GS samples was carried out by the Genetics Core Laboratory at the Edinburgh Clinical Research Facility, University of Edinburgh, Scotland, and was funded by the Medical Research Council UK and the Wellcome Trust (Wellcome Trust Strategic Award ‘Stratifying Resilience and Depression Longitudinally’ Reference 104036/Z/14/Z). DNA methylation profiling was also funded in part by the Wellcome Trust Investigator Award (Reference 220857/Z/20/Z). High-performance computing was supported by the Scientific Service Units (SSU) of IST Austria through resources provided by Scientific Computing (SciComp). MM, AD and YZ are funded by the European Union (ERC, INF2, project number 101161364). Views and opinions expressed are however those of the author(s) only and do not necessarily reflect those of the European Union or the European Research Council Executive Agency. Neither the European Union nor the granting authority can be held responsible for them. The work of AS and YZ was done while they were interns at ISTA.

## Author Contributions Statement

AD, MM and MRR conceived the study. AD, JB, MM and MRR designed the study. AD derived the model and the algorithm, with contributions from AS, JB, YZ, MM and MRR. JB, AD, and YZ prepared the data. JB, AD, YZ and AS wrote the software, and conducted the analyses. AL, AR and REM conducted the replication analyses. SV, AG, SA, AZ and MR were involved in the provision of data for replication analysis. HK, REM, MM and MRR provided study oversight. JB, AD, MM and MRR wrote the paper. All authors approved the final manuscript prior to submission.

## Data Availability

This project uses the UK Biobank data under project number 35520. UK Biobank genotypic and phenotypic data is available through a formal request at (http://www.ukbiobank.ac.uk). According to the terms of consent for Generation Scotland participants, access to data must be reviewed by the Generation Scotland Access Committee. Access requests can be made through the researcher portal (https://gsaccess.igc.ed.ac.uk) and normally take up to six weeks for approval. Further details can be found at https://genscot.ed.ac.uk/for-researchers/access/. All summary statistic estimates are released publicly on Dryad: https://doi.org/10.5061/dryad.9ghx3fg01. Source Data are provided with this paper.

## Code Availability

Source code for vampW, along with the code for reproducing the analyses in this paper, can be found at https://github.com/Information-and-learning-for-genomics/Time2EVAMP.

## Disclosure of the Use of Generative AI

AI generative tools were used for text grammar polishing and code debugging. All AI-generated content was reviewed and fact-checked by the authors prior to further development. The authors take full responsibility for the content.

## Supplementary information

### Supplementary Tables

**Supplementary Table 1:** Description of parameters varied in the simulation study. We simulate 24 artificial phenotypic outcomes covering a broad range of parameterizations, varying the variance explained by protein levels (*σ*_*G*_), the censoring ratio (*C*), the phenotype distribution, the ExpGamma parameter (*κ*), and the distribution of regression coefficients (***β***).

| Label | $\sigma_G^2$ | $C$ | Phenotype distribution | $\kappa$ | Prior $\beta$ distribution |
| --- | --- | --- | --- | --- | --- |
| a | 40% | 90% | Weibull | - | Normal |
| b | 80% | 90% | Weibull | - | Normal |
| c | 40% | 90% | ExpGamma | 0.8 | Normal |
| d | 80% | 90% | ExpGamma | 0.8 | Normal |
| e | 40% | 90% | ExpGamma | 1.2 | Normal |
| f | 80% | 90% | ExpGamma | 1.2 | Normal |
| g | 40% | 90% | Weibull | - | Laplace |
| h | 80% | 90% | Weibull | - | Laplace |
| i | 40% | 90% | ExpGamma | 0.8 | Laplace |
| j | 80% | 90% | ExpGamma | 0.8 | Laplace |
| k | 40% | 90% | ExpGamma | 1.2 | Laplace |
| l | 80% | 90% | ExpGamma | 1.2 | Laplace |
| m | 40% | 95% | Weibull | - | Normal |
| n | 80% | 95% | Weibull | - | Normal |
| o | 40% | 95% | ExpGamma | 0.8 | Normal |
| p | 80% | 95% | ExpGamma | 0.8 | Normal |
| q | 40% | 95% | ExpGamma | 1.2 | Normal |
| r | 80% | 95% | ExpGamma | 1.2 | Normal |
| s | 40% | 95% | Weibull | - | Laplace |
| t | 80% | 95% | Weibull | - | Laplace |
| u | 40% | 95% | ExpGamma | 0.8 | Laplace |
| v | 80% | 95% | ExpGamma | 0.8 | Laplace |
| w | 40% | 95% | ExpGamma | 1.2 | Laplace |
| x | 80% | 95% | ExpGamma | 1.2 | Laplace |

**Supplementary Table 2:** Number of significant protein findings per trait and method. vampW: PIP ≥ 0.95; CoxPH and CoxPH marginal: *p* ≤ 0.05/*M*; CoxPH LASSO and CoxPH Elastic Net (EN): *p* ≤ 0.05/*M*_stable_.

| Trait | vampW | CoxPH | CoxPH marginal | CoxPH LASSO | CoxPH EN |
| --- | --- | --- | --- | --- | --- |
| Alzheimer | 18 | 0 | 0 | 10 | 10 |
| Asthma | 0 | 12 | 1101 | 52 | 49 |
| COPD | 0 | 10 | 1153 | 32 | 41 |
| Cystitis | 0 | 1 | 4 | 7 | 8 |
| Death | 16 | 14 | 1367 | 34 | 34 |
| Depression | 69 | 9 | 857 | 45 | 40 |
| Endometriosis | 4 | 4 | 17 | 13 | 11 |
| Hypertension | 52 | 27 | 1826 | 94 | 94 |
| Inflammatory bowel disease | 0 | 6 | 305 | 34 | 35 |
| Ischemic heart disease | 19 | 10 | 956 | 37 | 46 |
| Ischemic stroke | 0 | 15 | 646 | 20 | 20 |
| Liver disease | 1 | 0 | 724 | 9 | 11 |
| Multiple sclerosis | 0 | 4 | 65 | 26 | 25 |
| Parkinson | 11 | 5 | 13 | 38 | 36 |
| Rheumatoid arthritis | 5 | 18 | 859 | 16 | 21 |
| Systemic lupus erythematosus | 0 | 2 | 713 | 10 | 19 |
| Type 2 diabetes | 13 | 18 | 1575 | 62 | 62 |
| Vascular dementia | 0 | 0 | 0 | 4 | 0 |
| Breast cancer | 2 | 12 | 80 | 28 | 35 |
| Brain/CNS cancer | 0 | 0 | 0 | 6 | 4 |
| Colorectal cancer | 1 | 1 | 8 | 8 | 6 |
| Gynecological cancer | 1 | 1 | 11 | 9 | 9 |
| Lung cancer | 2 | 0 | 200 | 15 | 15 |
| Prostate cancer | 5 | 16 | 5 | 20 | 15 |
| Total | 219 | 185 | 12485 | 629 | 646 |

**Supplementary Table 3:** Overlap between the top-*x* findings of each pair of methods, summed over traits. For each trait, both lists are ranked by significance (vampW by PIP, Cox models by *p*-value) and truncated to *x* = min(*n*_1_, *n*_2_), the smaller number of findings of the two methods. Cells show overlap / Σ*x* The diagonal gives the total number of findings of each method.

|  | vampW | CoxPH | CoxPH marginal | CoxPH LASSO | CoxPH EN |
| --- | --- | --- | --- | --- | --- |
| vampW | 219 | 45 / 96 | 37 / 201 | 85 / 187 | 90 / 182 |
| CoxPH | 45 / 96 | 185 | 28 / 174 | 78 / 183 | 81 / 184 |
| CoxPH marginal | 37 / 201 | 28 / 174 | 12485 | 112 / 566 | 110 / 595 |
| CoxPH LASSO | 85 / 187 | 78 / 183 | 112 / 566 | 629 | 510 / 603 |
| CoxPH EN | 90 / 182 | 81 / 184 | 110 / 595 | 510 / 603 | 646 |

**Supplementary Table 4:** Number of significant (trait, protein) findings shared by each pair of methods, summed over traits. The diagonal gives the total number of findings of each method. Traits without a result for one of the two methods are excluded from that pair.

|  | vampW | CoxPH | CoxPH marginal | CoxPH LASSO | CoxPH EN |
| --- | --- | --- | --- | --- | --- |
| vampW | 219 | 61 | 138 | 110 | 112 |
| CoxPH | 61 | 185 | 110 | 112 | 116 |
| CoxPH marginal | 138 | 110 | 12485 | 385 | 407 |
| CoxPH LASSO | 110 | 112 | 385 | 629 | 532 |
| CoxPH EN | 112 | 116 | 407 | 532 | 646 |

**Supplementary Table 5:** Description of traits under investigation from the UK Biobank health records. For cancer traits, cases are defined by the presence of disease-specific ICD-10 codes (field 40006) or ICD-9 codes (field 40013) across all instances. For the non-cancer traits, we analyze time-to-event derived from the first reported occurrence of each disease (category 1712). The column ‘Cases’ reports the total number of individuals who experienced the event (non-censored observations) within the study cohort.

| Abbreviation | Phenotype | UK Biobank code | Cases |
| --- | --- | --- | --- |
| CNS | Brain/CNS cancer | 100092 (cancer register) | 115 |
| MS | Multiple sclerosis | 131042 | 508 |
| MD | Major depression | 130894 OR 130896 | 7,128 |
| SLE | Systemic lupus erythematosus | 131894 | 579 |
| ENDO | Endometriosis | 132122 | 1,031 |
| VD | Vascular dementia | 130838 | 297 |
| GC | Gynecological cancer | 100092 (cancer register) | 664 |
| ALS | Amyotrophic lateral sclerosis | 42028 | 508 |
| IBD | Inflammatory bowel disease | 131626 OR 131628 | 1,014 |
| LC | Lung cancer | 100092 (cancer register) | 598 |
| LD | Liver disease | 131658 | 268 |
| AD | Alzheimer’s dementia | 130836 | 546 |
| CD | Colorectal cancer | 100092 (cancer register) | 988 |
| CYS | Cystitis | 132054 | 2,169 |
| RA | Rheumatoid arthritis | 131850 | 1,556 |
| PD | Parkinson’s disease | 131022 | 963 |
| IS | Ischemic stroke | 131368 | 1,255 |
| BC | Breast cancer | 100092 (cancer register) | 1,983 |
| PC | Prostate cancer | 100092 (cancer register) | 1,671 |
| COPD | COPD | 131492 | 3,394 |
| T2D | Type 2 diabetes | 130708 (T1D 130706 absent) | 4,760 |
| IHD | Ischemic heart disease | 131306 | 6,438 |
| DEATH | Death | 100093 (death register) | 5,765 |
| HTN | Hypertension | 131286 | 22,408 |

**Supplementary Table 6:**
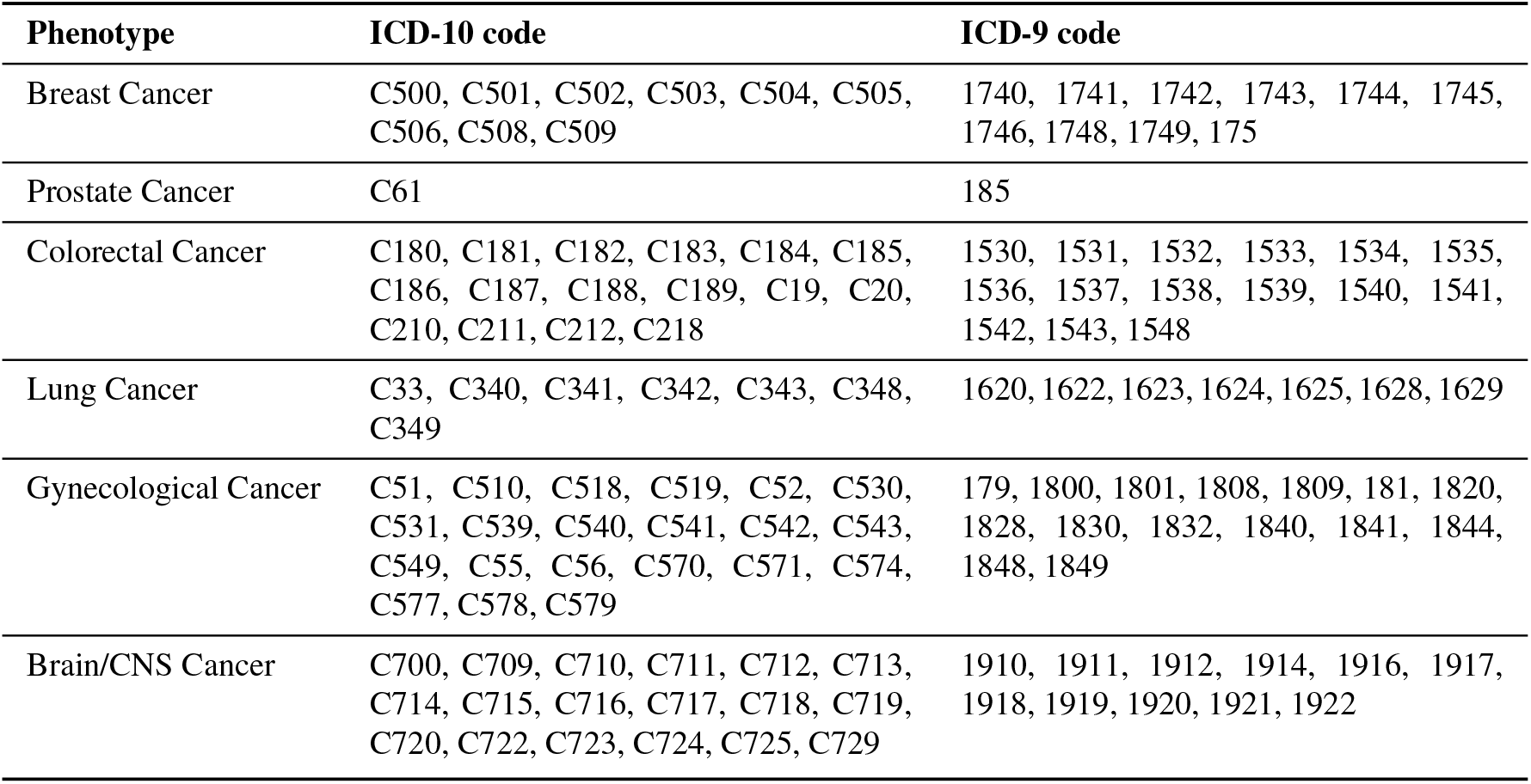
ICD-9 and ICD-10 codes used to define cancer phenotypes in the UK Biobank. Cases were identified by matching these codes against the UK Biobank cancer registry records (ICD-10 field 40006 and ICD-9 field 40013) to determine the age at diagnosis.

| Phenotype | ICD-10 code | ICD-9 code |
| --- | --- | --- |
| Breast Cancer | C500, C501, C502, C503, C504, C505, C506, C508, C509 | 1740, 1741, 1742, 1743, 1744, 1745, 1746, 1748, 1749, 175 |
| Prostate Cancer | C61 | 185 |
| Colorectal Cancer | C180, C181, C182, C183, C184, C185, C186, C187, C188, C189, C19, C20, C210, C211, C212, C218 | 1530, 1531, 1532, 1533, 1534, 1535, 1536, 1537, 1538, 1539, 1540, 1541, 1542, 1543, 1548 |
| Lung Cancer | C33, C340, C341, C342, C343, C348, C349 | 1620, 1622, 1623, 1624, 1625, 1628, 1629 |
| Gynecological Cancer | C51, C510, C518, C519, C52, C530, C531, C539, C540, C541, C542, C543, C549, C55, C56, C570, C571, C574, C577, C578, C579 | 179, 1800, 1801, 1808, 1809, 181, 1820, 1828, 1830, 1832, 1840, 1841, 1844, 1848, 1849 |
| Brain/CNS Cancer | C700, C709, C710, C711, C712, C713, C714, C715, C716, C717, C718, C719, C720, C722, C723, C724, C725, C729 | 1910, 1911, 1912, 1914, 1916, 1917, 1918, 1919, 1920, 1921, 1922 |

**Supplementary Table 7:** Prior initialization configurations for vampW. For analyzing real UK Biobank traits, we use several vampW initializations, varying the damping factor (*ρ*), the initial Weibull shape parameter (*α*), and the prior variances and probabilities.

| Label | $\rho$ | Initial $\alpha$ | Variances | Probabilities | Adaptive $\rho$ |
| --- | --- | --- | --- | --- | --- |
| a | 0.3 | 3 | 0, 0.01 | 0.7, 0.3 | No |
| b | 0.5 | 3 | 0, 0.01 | 0.7, 0.3 | No |
| c | 0.5 | 3 | 0, 0.01 | 0.7, 0.3 | Yes |
| d | 0.5 | 3 | 0, 1 | 0.7, 0.3 | Yes |
| e | 0.6 | 3 | 0, 1 | 0.7, 0.3 | No |
| f | 0.8 | 3 | 0, 0.01 | 0.7, 0.3 | No |
| g | 0.8 | 3 | 0, 0.01 | 0.7, 0.3 | Yes |
| h | 1.0 | 3 | 0, 1 | 0.7, 0.3 | No |

**Supplementary Table 8:** Initialization settings used per UK Biobank trait. We use different prior initializations, described in Supplementary Table 7, across the various traits. The ‘Initialization Label’ links each specific disease outcome to the corresponding hyperparameter configuration defined in Supplementary Table 7, ensuring reproducibility of the analysis settings.

| Trait | Initialization label |
| --- | --- |
| Alzheimer | b |
| Asthma | b |
| COPD | g |
| Cystitis | g |
| Death | h |
| Depression | c |
| Endometriosis | e |
| Hypertension | d |
| Inflammatory Bowel Disease | a |
| Ischemic Heart Disease | f |
| Ischemic Stroke | g |
| Liver Disease | g |
| Multiple Sclerosis | f |
| Parkinson | c |
| Rheumatoid Arthritis | g |
| Schizophrenia | g |
| Systemic Lupus Erythematosus | f |
| Type 2 Diabetes (T2D) | b |
| Vascular Dementia | f |
| Breast cancer | g |
| Brain CNS cancer | g |
| Colorectal cancer | g |
| Gynecological cancer | g |
| Lung cancer | g |
| Prostate cancer | g |

### Supplementary Figures

**Supplementary Figure 1:**
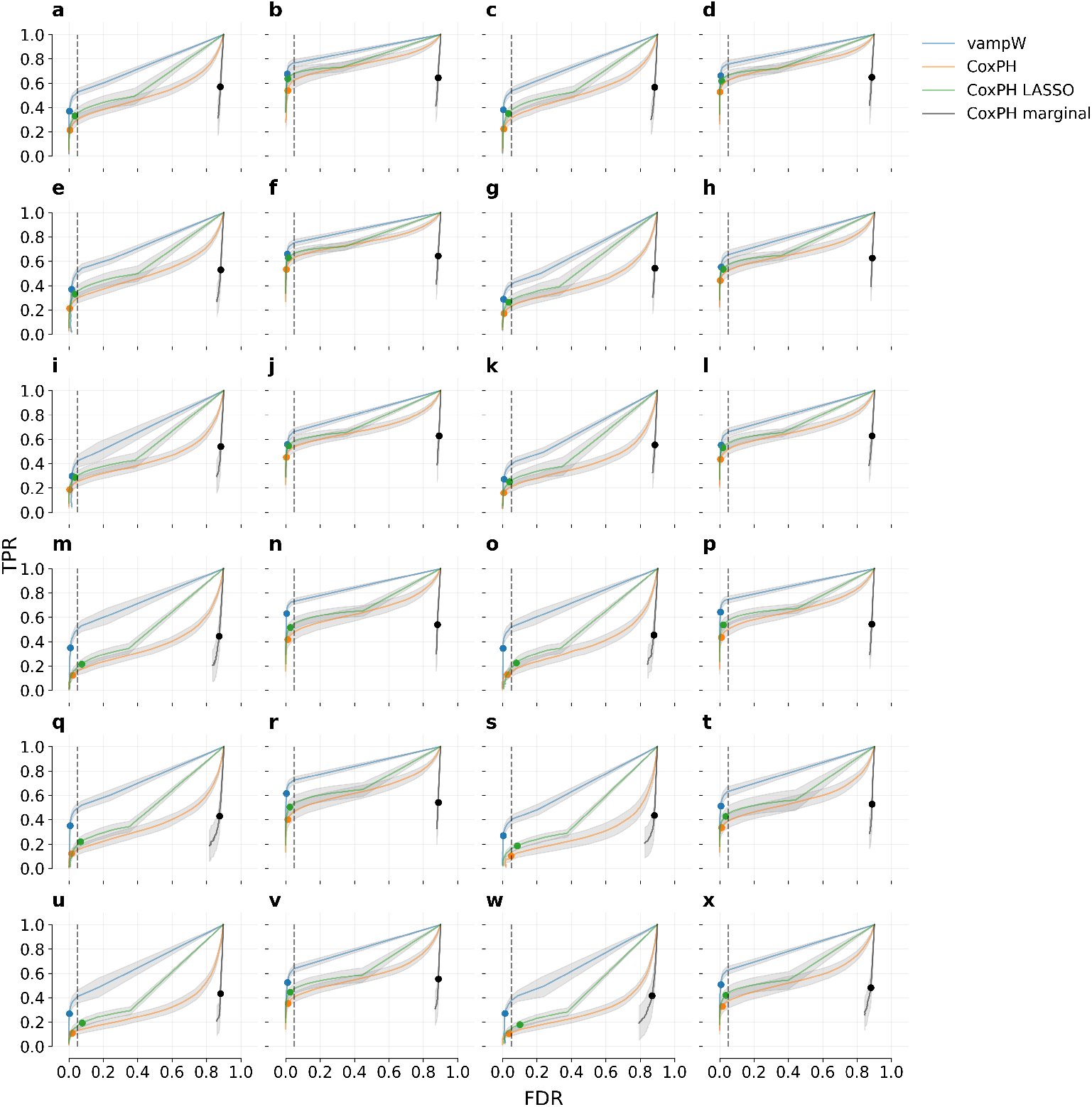
Simulation study of variable selection performance using protein data from the UK Biobank. We simulate artificial phenotypic outcomes under 24 scenarios, corresponding to the parameterizations described in Supplementary Table 1, varying the proportion of variance explained, level of censoring, phenotype distribution, and distribution of regression coefficients. We compare the variable selection performance of vampW against several benchmarking methods (CoxPH, CoxPH LASSO and CoxPH marginal) in terms of the relationship between the false discovery rate (FDR) and true positive rate (TPR). For vampW, we use posterior inclusion probabilities (PIPs), with colored dots indicating a 0.95 threshold. For the other methods, p-value testing across multiple thresholds is employed, with colored dots indicating a Bonferroni-corrected 0.05 threshold. vampW maintains a calibrated FDR and consistently outperforms CoxPH-based models in TPR across all simulation scenarios, including those involving model misspecification. The error bars (gray shaded areas) represent the standard deviation across 50 simulation replicates.

**Supplementary Figure 2:**
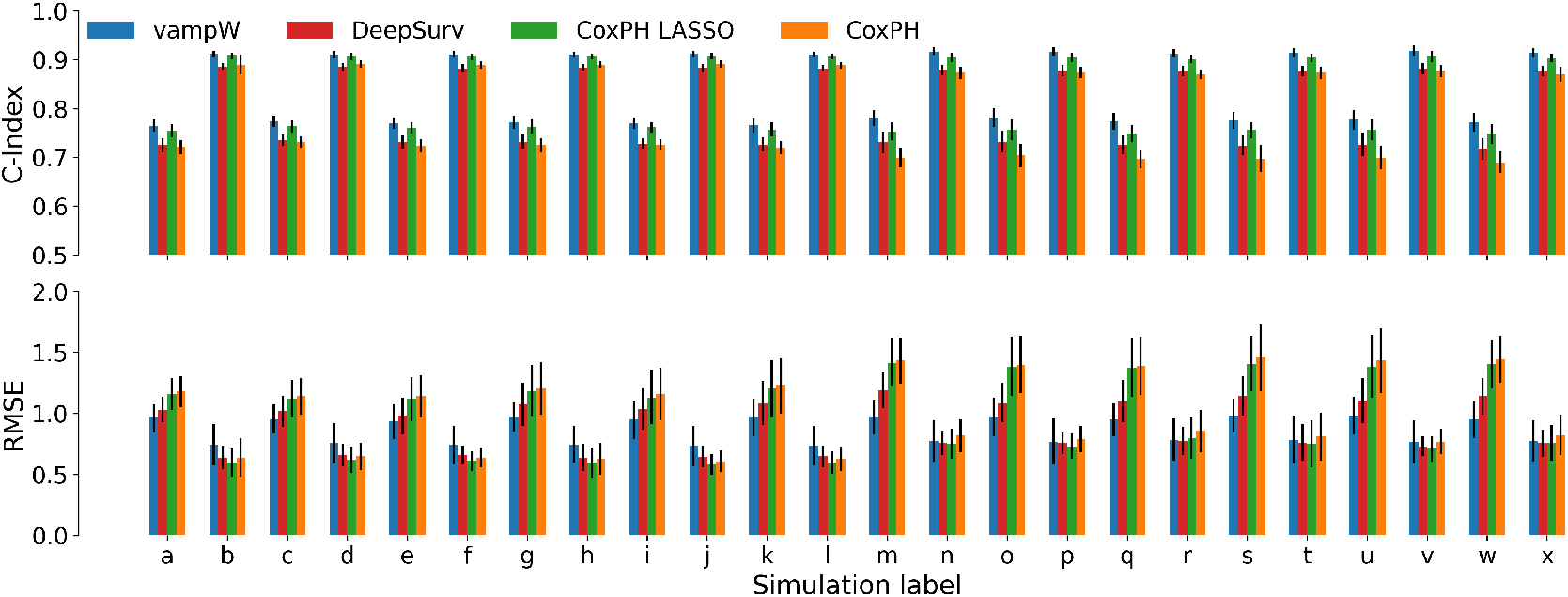
Simulation study of out-of-sample prediction performance using protein data from the UK Biobank. We evaluate predictive performance for time of onset across 24 simulated phenotypic outcomes corresponding to the parameterizations in Supplementary Table 1. We compare vampW with CoxPH, CoxPH LASSO, and DeepSurv in terms of concordance index (C-index) and root mean squared error (RMSE). The RMSE is evaluated in the logarithmic domain, assuming standardized outcomes with zero mean and unit variance. Across all simulated scenarios, vampW consistently demonstrates a clear improvement in the C-index when compared to both DeepSurv and the standard CoxPH model. When compared to CoxPH LASSO, the point estimate of C-index values for vampW improves, although it remains within overlapping error bars across all scenarios. In terms of RMSE, vampW strictly outperforms the standard CoxPH model in six scenarios and improves upon CoxPH LASSO in five scenarios. In all remaining comparisons, the predictive performance of the evaluated methods falls within each other’s error margins. The reported error bars represent the standard deviation across 50 simulation replicates (black lines at the top of the colored bars).

**Supplementary Figure 3:**
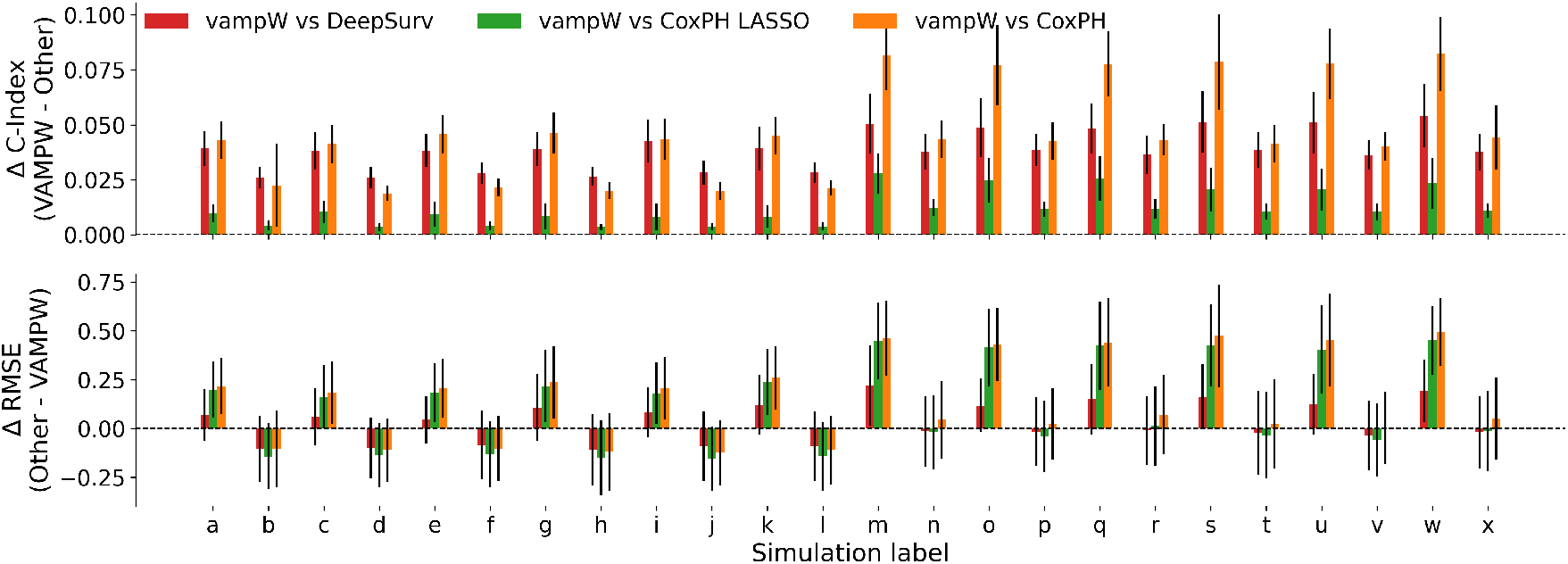
Paired differences in out-of-sample prediction performance across simulated phenotypic outcomes. We evaluate the predictive performance of vampW compared to DeepSurv, CoxPH LASSO, and the standard CoxPH model across 24 simulated scenarios (corresponding to the parameterizations in Supplementary Table 1). The top panel displays the paired difference in concordance index (Δ C-index), calculated as vampW minus the competing method. The bottom panel shows the paired difference in root mean squared error (Δ RMSE), calculated as the competing method minus vampW. By this formulation, positive values in both panels (bars extending above the dashed zero line) indicate superior predictive performance by vampW. The RMSE is evaluated in the logarithmic domain, assuming standardized outcomes with zero mean and unit variance. The reported error bars represent the standard deviation of the paired differences across 50 simulation replicates (black lines at the top or bottom of the colored bars). For the C-index, the paired differences, inclusive of error bars, never cross the zero threshold, indicating that vampW consistently outperforms the other methods across all scenarios. In terms of RMSE, vampW achieves performance that is either on par with or superior to the competing methods.

**Supplementary Figure 4:**
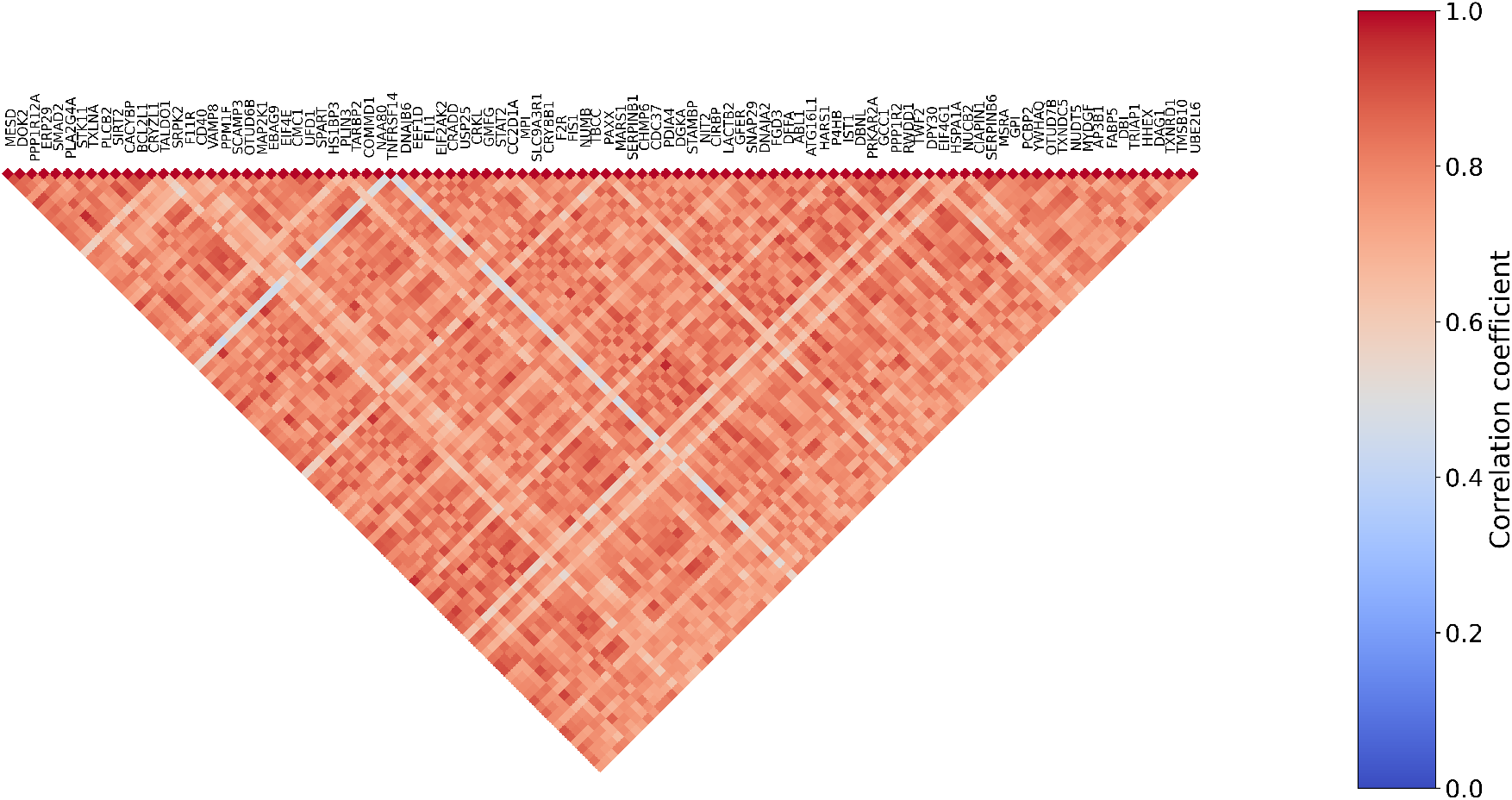
Correlation structure of the protein measures in the UK Biobank. We calculate Pearson product-moment correlation coefficients for all proteins, not-adjusted for covariates, and select the top 100 most correlated ones. We plot the correlation structure along with corresponding protein-coding genes.

**Supplementary Figure 5:**
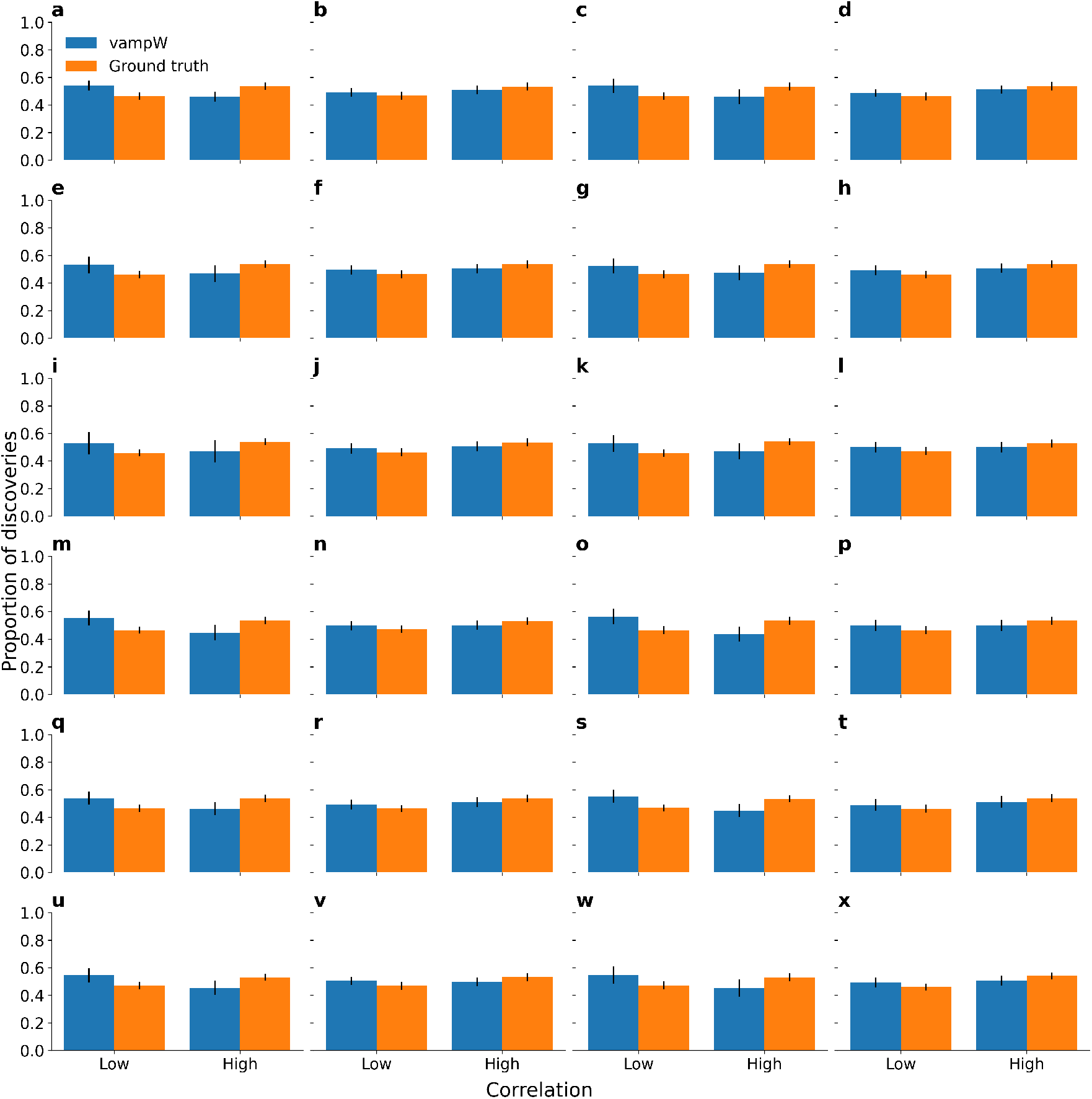
Protein annotation based on correlation structure. For 24 simulation scenarios, described in Supplementary Table 1, we visualize the relative proportion of highly and low-correlated true positive discoveries with respect to all true positive discoveries (blue), and compare this to the proportion of highly and low-correlated causal proteins with respect to all causal proteins (orange). If the correlation of a particular protein with any other protein is greater than 0.5 (absolute value of Pearson correlation coefficient), it is categorized as a highly correlated protein. The remaining proteins are categorized as low-correlated. The black bars show 1 standard error over 50 simulation replicas. The results suggest that vampW detects causal signals even for proteins with correlations above the 0.5 threshold.

**Supplementary Figure 6:**
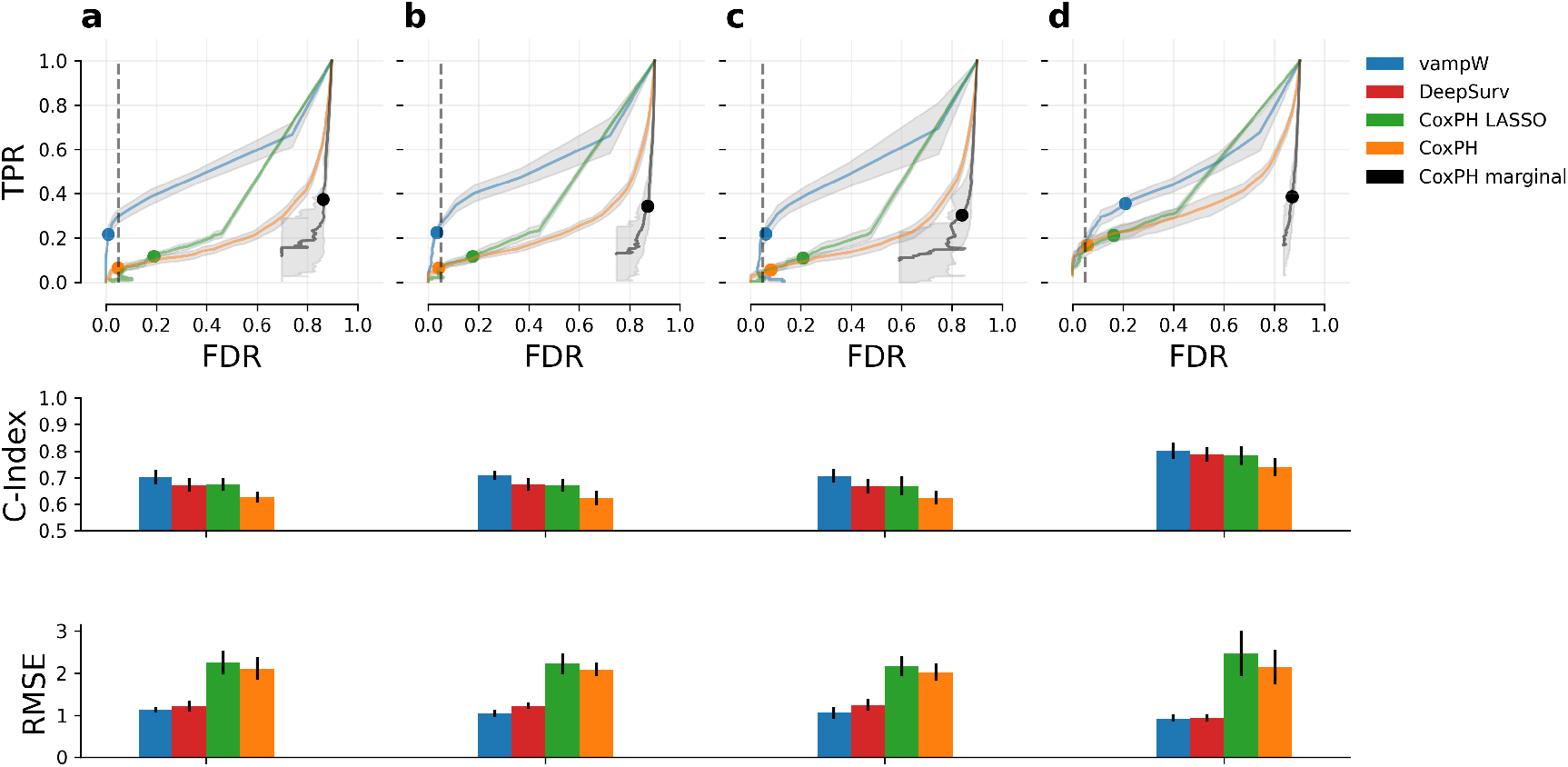
Including non-linear effects in simulation study of variable selection and out-of-sample prediction performance. The four scenarios, a–d, correspond to the simulation parameterizations (*m, o, q, t*) from Supplementary Table 1, covering model misspecifications and low and high signal-to-noise ratios. In addition, we extend the simulations by assigning 50% of the signal to linear effects and 50% to quadratic effects, while retaining the same causal positions. We also extend the grid parameter space for DeepSurv optimization by increasing the number of hidden layers to accommodate more complex effects. vampW achieves performance within DeepServ’s error bars in three of four scenarios in C-index, while performing better on average in all cases. In RMSE, vampW performs best in three out of four scenarios. The error bars represent the standard deviation across 10 simulation replicates.

**Supplementary Figure 7:**
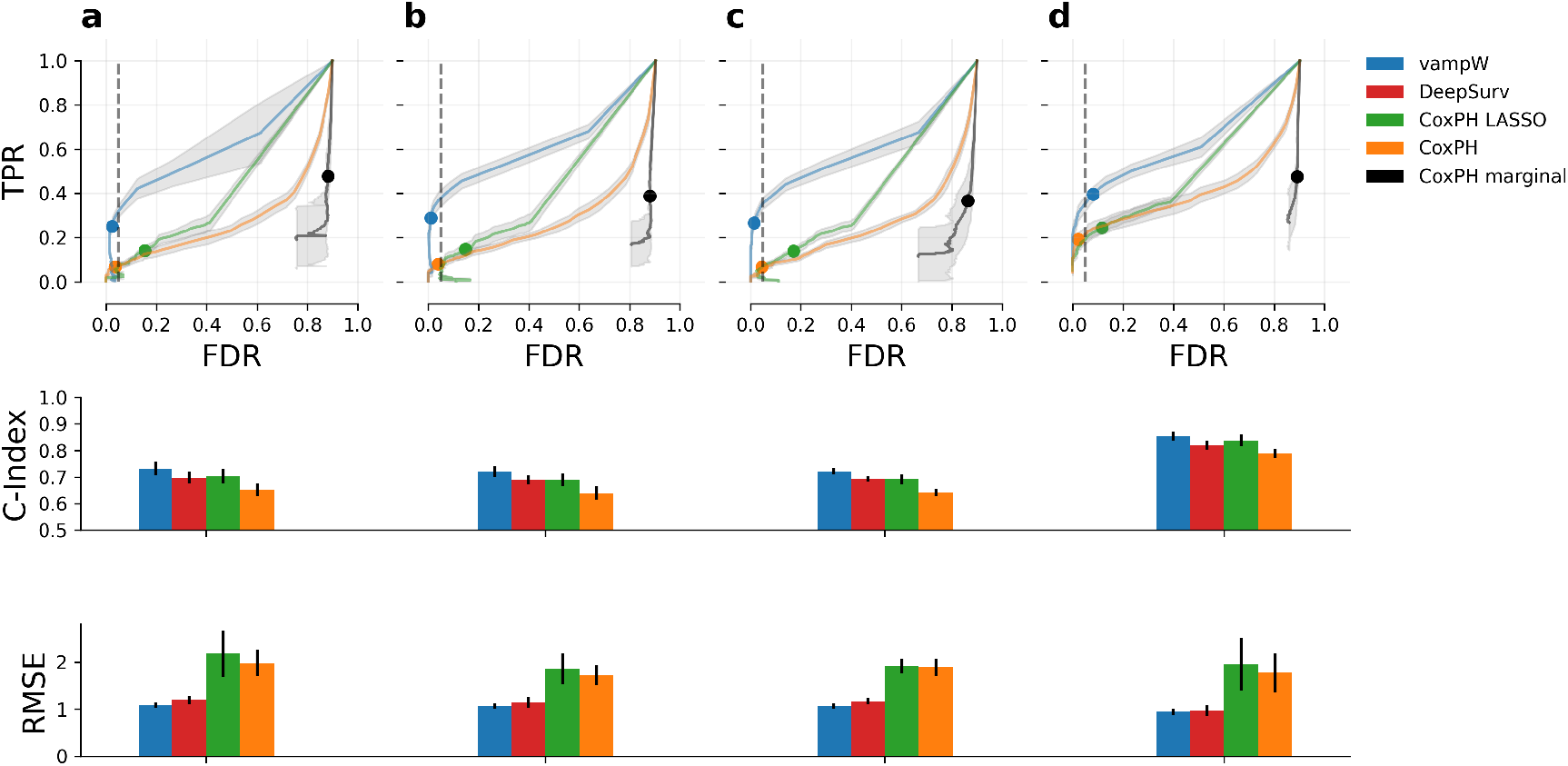
Including interaction effects in a simulation study of variable selection and out-of-sample prediction performance. The four scenarios, a–d, correspond to the simulation parameterizations (*m, o, q, t*) from Supplementary Table 1, covering model misspecifications and low and high signal-to-noise ratios. In addition, we extend the simulations by assigning 20% of the overall signal to pairwise protein–protein interactions, represented as a multiplication term of two interacting proteins selected from the STRING database. We also extend the grid parameter space for DeepSurv optimization by increasing the number of hidden layers to accommodate more complex effects. On average, vampW performs better than competing methods in C-index. In RMSE, vampW and Deepsurv fall within the error bars, particularly in column d, where the signal-to-noise ratio is 80%. The error bars represent the standard deviation across 9 simulation replicates (we exclude one outlier, causing numerical instability).

**Supplementary Figure 8:**
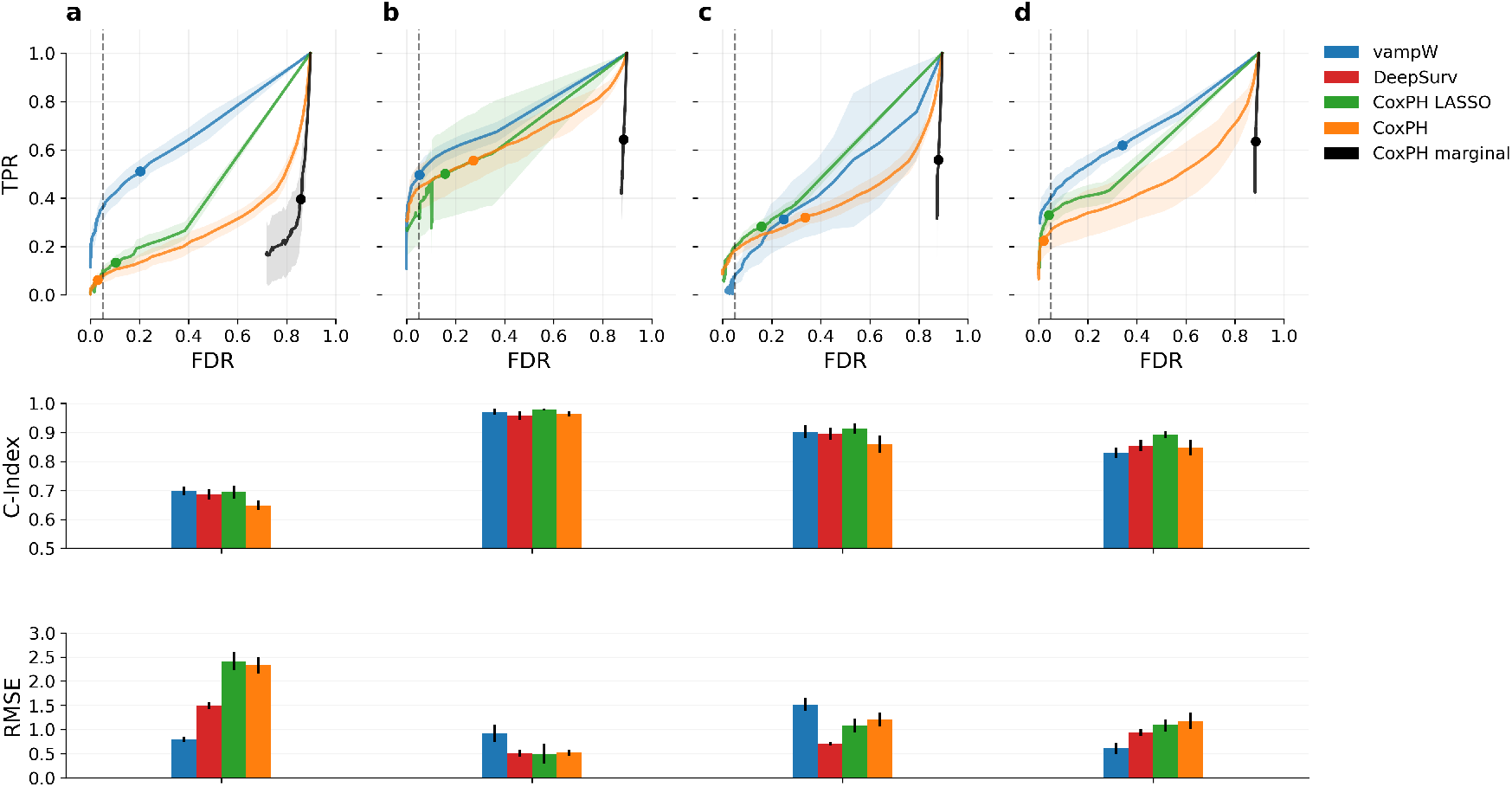
Including competing risk effects in a simulation study of variable selection and out-of-sample prediction performance. We further extend the simulation study by generating two event times, one for the primary event and one for the censoring process under the same distribution (Weibull or ExpGamma), while causal variants are drawn jointly from a bivariate distribution including correlation. We define observed time as the minimum of the two times, with the event indicator set to one when the primary event occurs first and zero when the competing event occurs first. The four scenarios, a–d, correspond to the simulation parameterizations (*m, o, q, t*) from Supplementary Table 1. vampW outperforms other methods in setups following the Weibull distribution (a,d) and loses in ExpGamma models (b,c) in RMSE while staying competitive in C-index. The error bars represent the standard deviation across 10 simulation replicates.

**Supplementary Figure 9:**
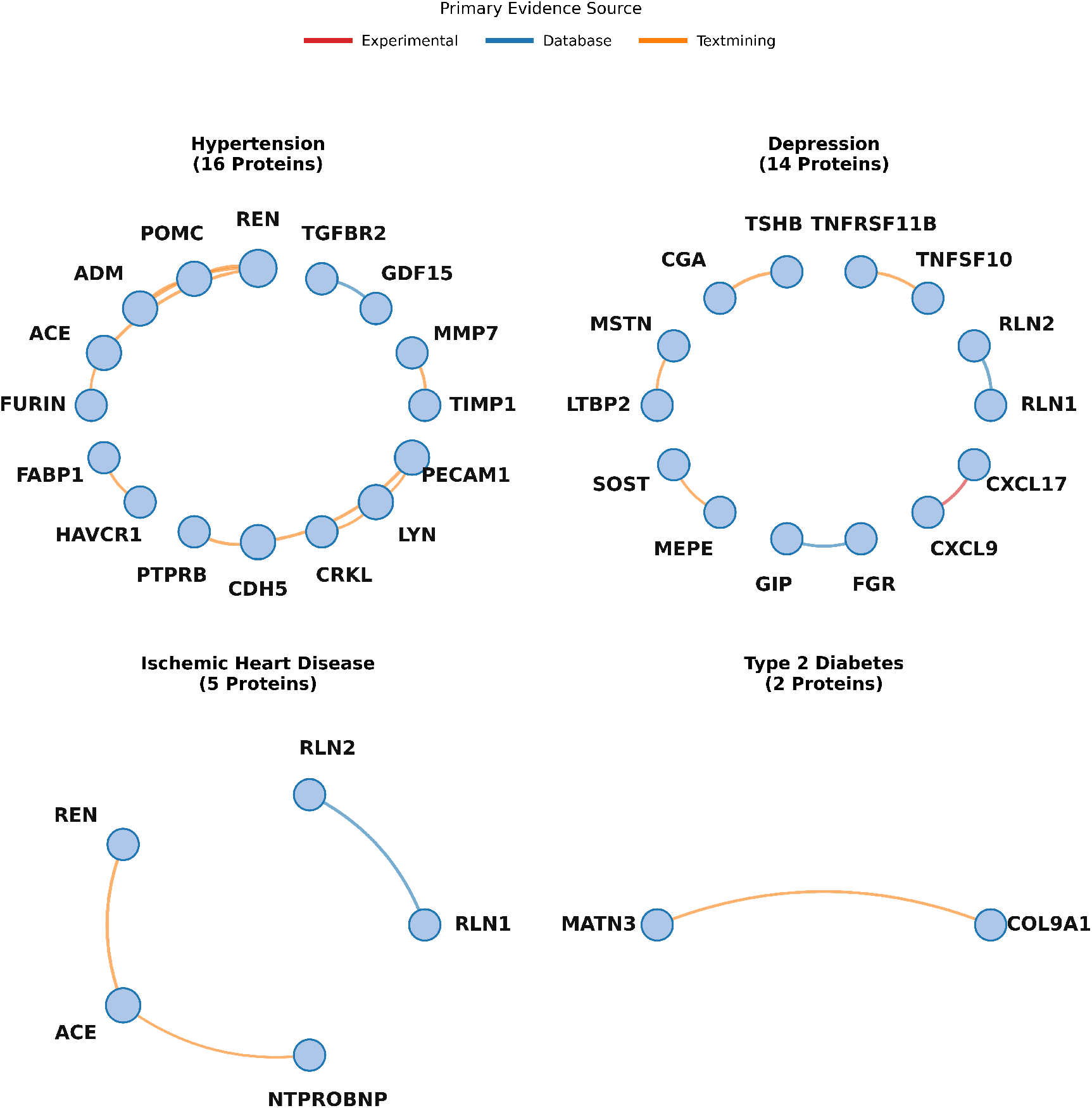
Protein-protein interaction networks for trait-associated proteins. Network visualizations of significant proteins associated with Hypertension, Depression, Ischemic Heart Disease, and Type 2 Diabetes. Nodes represent individual proteins identified in the analysis. Edges denote known interactions retrieved from the STRING database (version 12.0) [45], filtered for high-confidence connections (combined score ≥ 800). Edge colors indicate the primary source of evidence supporting the interaction: experimental validation (red), curated databases (blue), or automated text mining (orange). Node sizes are proportionally scaled by their degree (number of connections) within the local trait network. Proteins without high-confidence interactions in the selected sub-network are excluded from the visualization.

**Supplementary Figure 10:**
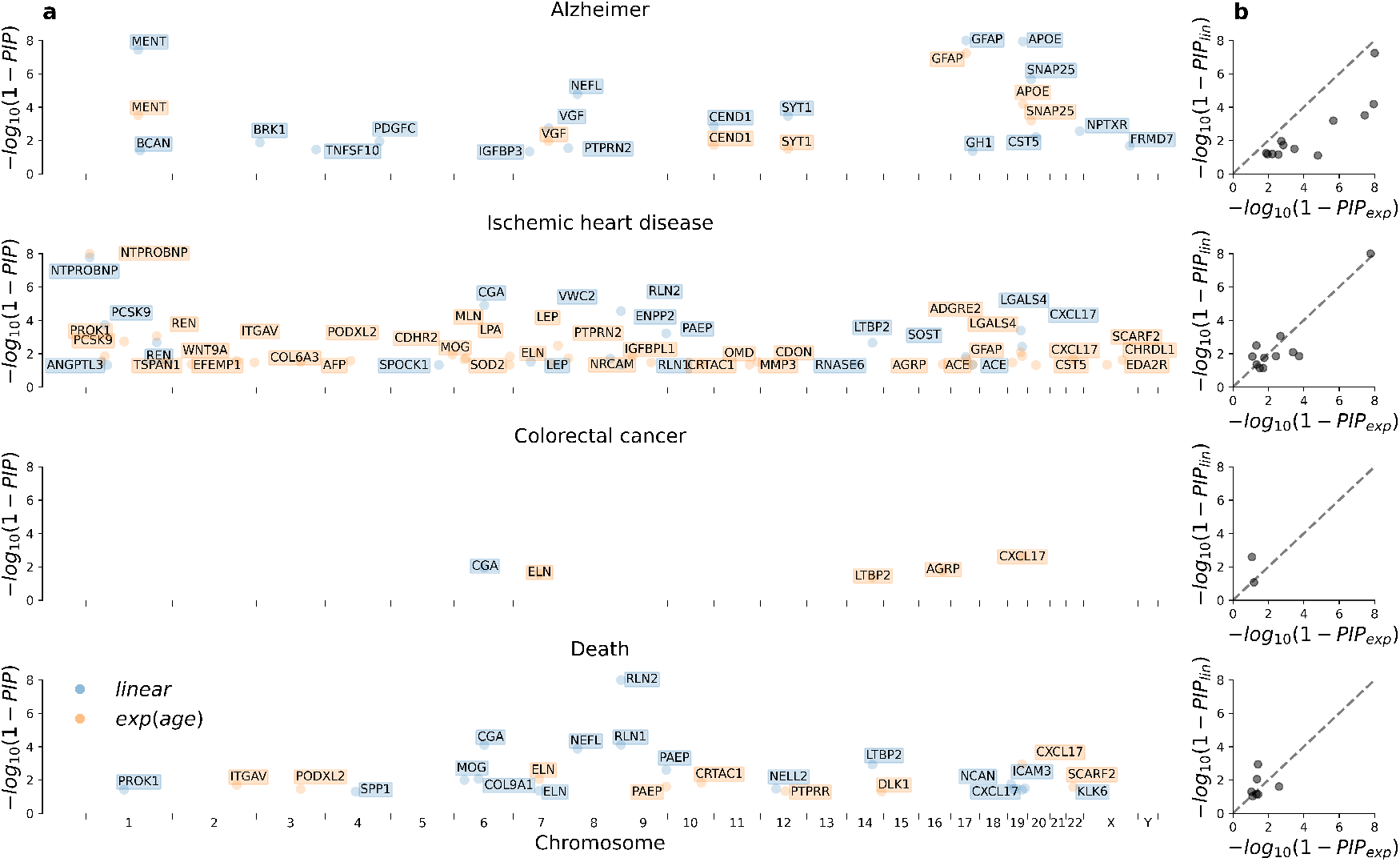
Adjusting for non-linear effects of age on protein levels. In addition to the standard linear adjustment (blue) for the age covariate, we evaluate the vampW model using protein data adjusted for exponential age effects (orange). In column (a), we show significant protein-coding genes at their corresponding DNA positions that pass a PIP threshold of 0.95 for four representative outcomes, namely Alzheimer’s disease, ischemic heart disease, colorectal cancer, and death. In column (b), we show the relationship between PIPs passing a threshold of 0.9 across the two models, where PIP_*lin*_ and PIP_*exp*_ denote the standard linear and exponential age adjustments, respectively. Notably, the gene *CGA*, previously reported as age-associated in recent work, disappears from the discoveries when considering non-linear effects of age on protein levels.

**Supplementary Figure 11:**
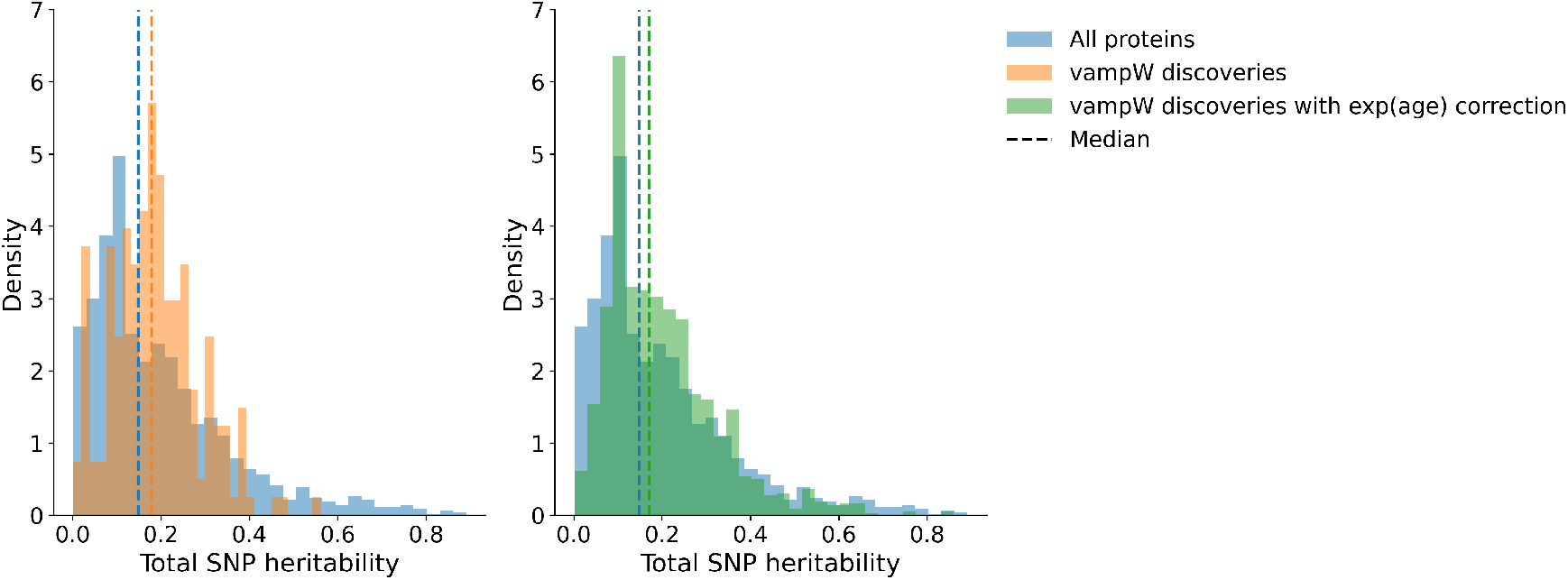
Distribution of total SNP-based heritability of proteins with pQTLs. We plot the distributions of total SNP heritability for 2,430 proteins reported in the study [3], including all proteins with pQTLs (blue) and the corresponding subset overlapping with vampW discoveries, with (green) and without (orange) exponential age correction. The vampW discoveries exhibit average heritability of 19.9% and 18.7% for models with and without exponential age correction, respectively, compared to a mean heritability of 19.5% across all proteins with pQTLs.

**Supplementary Figure 12:**
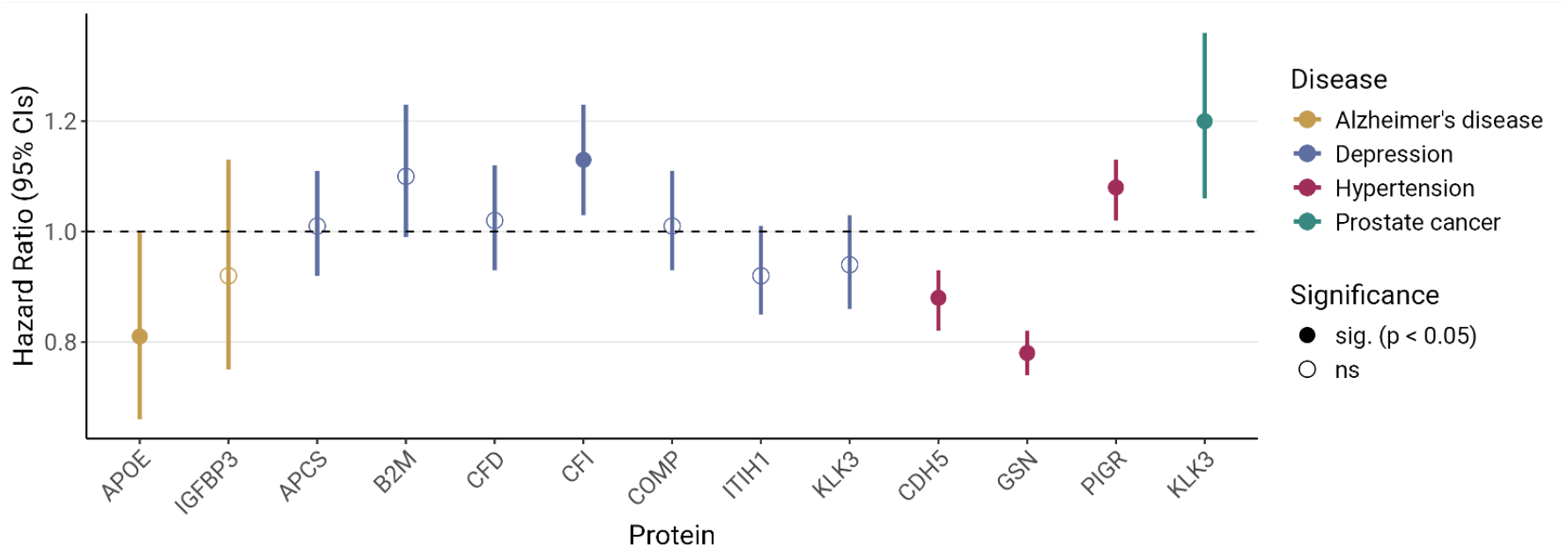
Replication in Generation Scotland mass spectrometry proteomics data. We analyze 13 protein–disease vampW associations from a model without exponential age correction, for which we find overlap in the Generation Scotland proteomics data. Specifically, we observe overlap for Alzheimer’s disease (yellow), depression (blue), hypertension (red), and prostate cancer (green), for which we run marginal Cox proportional hazards models for incident cases, adjusted for age and sex. We plot the corresponding hazard ratios with 95% confidence intervals (vertical lines) and indicate whether the p-value is below the 0.05 threshold (solid points) or not significant (ns). We find six replications without Bonferroni correction at the 5% threshold and four replications after Bonferroni correction at the 10% threshold.

**Supplementary Figure 13:**
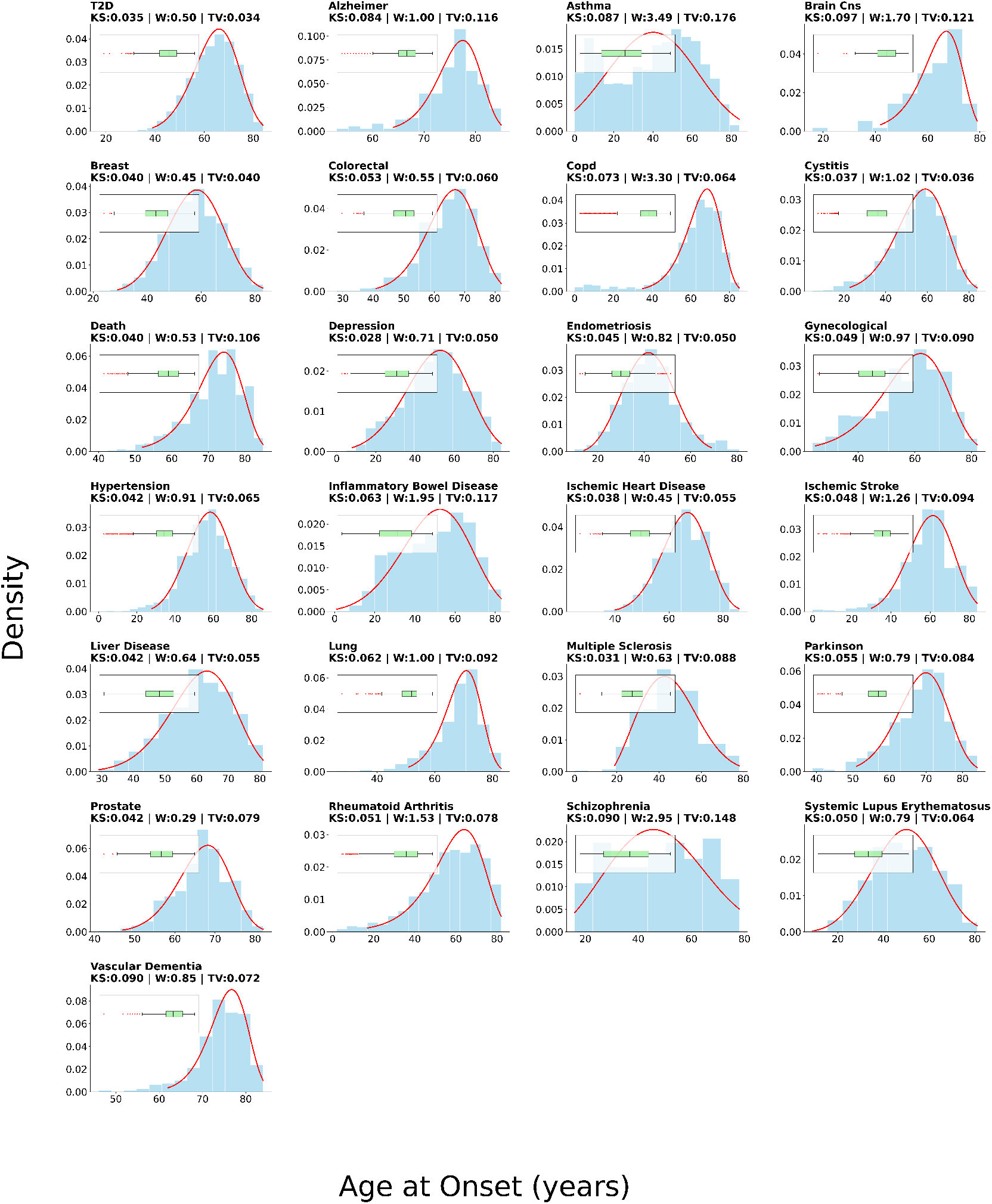
Analysis of age-at-onset distributions for UK Biobank traits. The histograms display the empirical distribution of age-at-onset for cases across various phenotypes in the training set. The blue bars represent the density of observed onset ages, while the solid red line indicates the fitted Weibull probability density function (PDF). Outliers were excluded from the fitting process using the Interquartile Range (IQR) method. The title of each subplot provides goodness-of-fit metrics, including the Kolmogorov-Smirnov statistic (KS), Wasserstein distance (W), and Total Variation distance (TV).

**Supplementary Figure 14:**
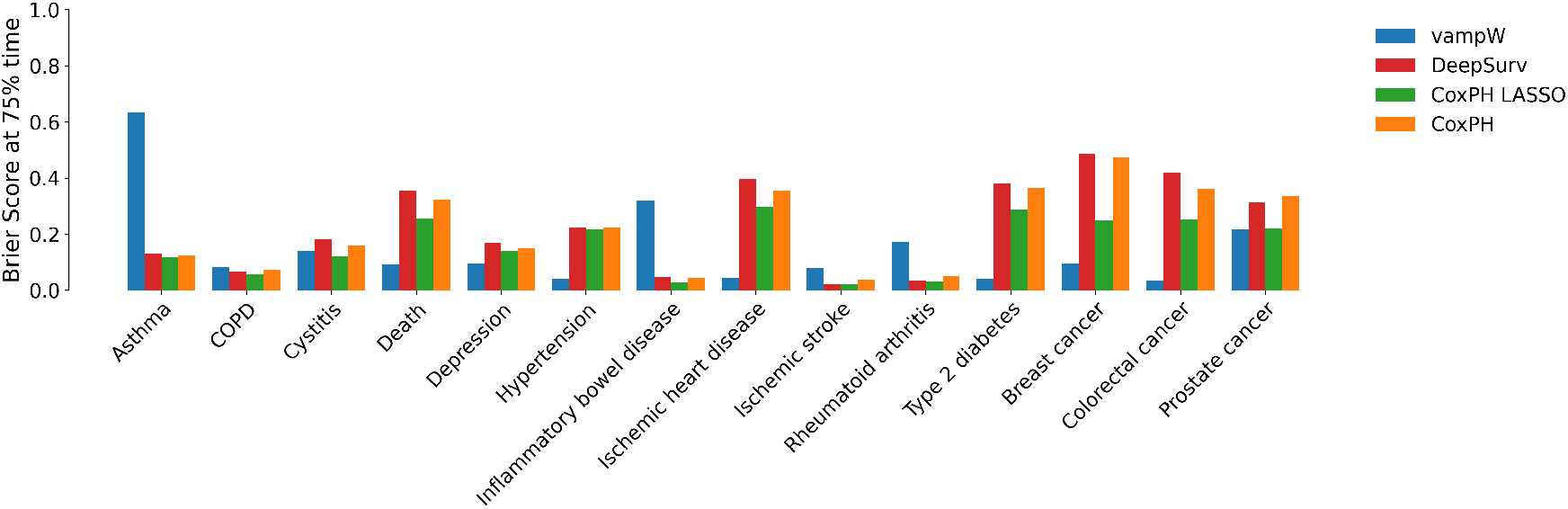
Brier Score comparison across methods evaluated at the 75% time point. We calculate the Brier Score (BS), a standard metric for evaluating the out-of-sample performance of time-to-event predictions. We report the BS value at the time corresponding to 75% of the time span within the observed time period. Among 14 UK Biobank outcomes with sufficient sample size for out-of-sample evaluation, vampW outperforms the other benchmarking methods in 8 cases. Based on Supplementary Figure 13, the Asthma distribution by age has one of the least Weibull-like shapes, which may drive the high Brier Score, as risk increases only modestly with age.

**Supplementary Figure 15:**
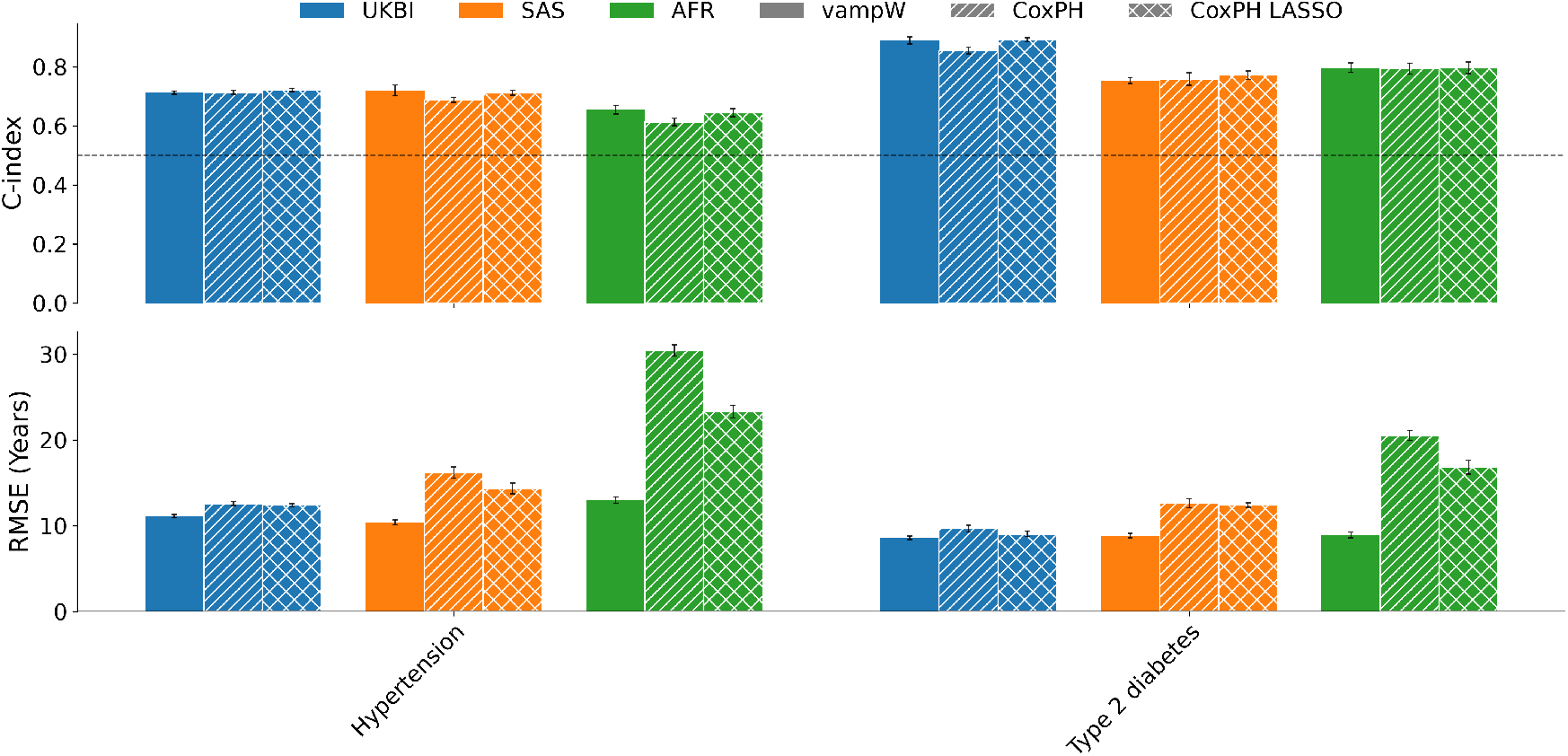
Out-of-sample prediction performance across distinct genetic ancestries. We evaluate the predictive performance of vampW (solid bars), the standard CoxPH model (diagonal hatch lines), and CoxPH LASSO (cross hatch lines) across Hypertension and Type 2 diabetes traits from the UK Biobank. Results are stratified by genetic ancestry: British or Irish (UKBI, blue), South Asian (SAS, orange), and African (AFR, green). Performance is measured using the concordance index (C-index, top panel) and root mean squared error (RMSE in years, bottom panel). Phenotypic traits were included only if at least one minority ancestry group (SAS or AFR) contained a minimum of 200 total cases, ensuring that the 50% data splits retained at least 100 cases per split. The reported error bars represent the standard deviation across 10 splits, with each split evaluating a random 50% subsample of the dataset. We note that polygenic risk scores created by vampW are transferable to minority populations in the cohort; vampW exhibits similar C-index performance to Cox LASSO and outperforms the joint standard Cox model, while outperforming both competitor models in terms of the RMSE metric.

**Supplementary Figure 16:**
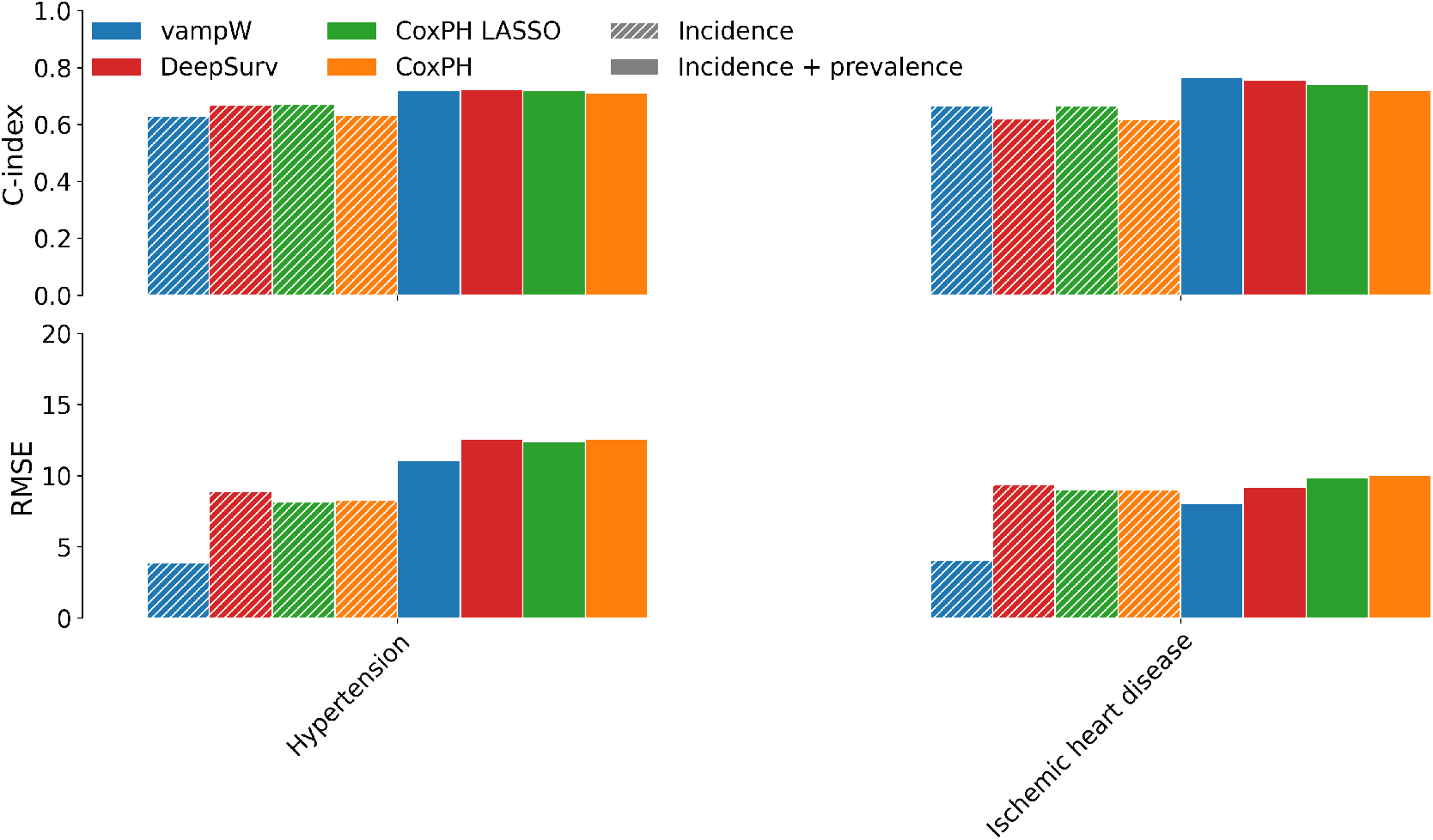
Time-to-event analysis on incidence cases in comparison to mixed prevalence and incident age-at-onset. We perform an incident-cases analysis for hypertension and ischemic heart disease, which have sufficient sample sizes. In the incidence analysis, vampW stays competitive to other methods in C-index while outperforming significantly in RMSE. vampW’s C-index performance drops by 12.61% for hypertension and 12.97% for ischemic heart disease.

**Supplementary Figure 17:**
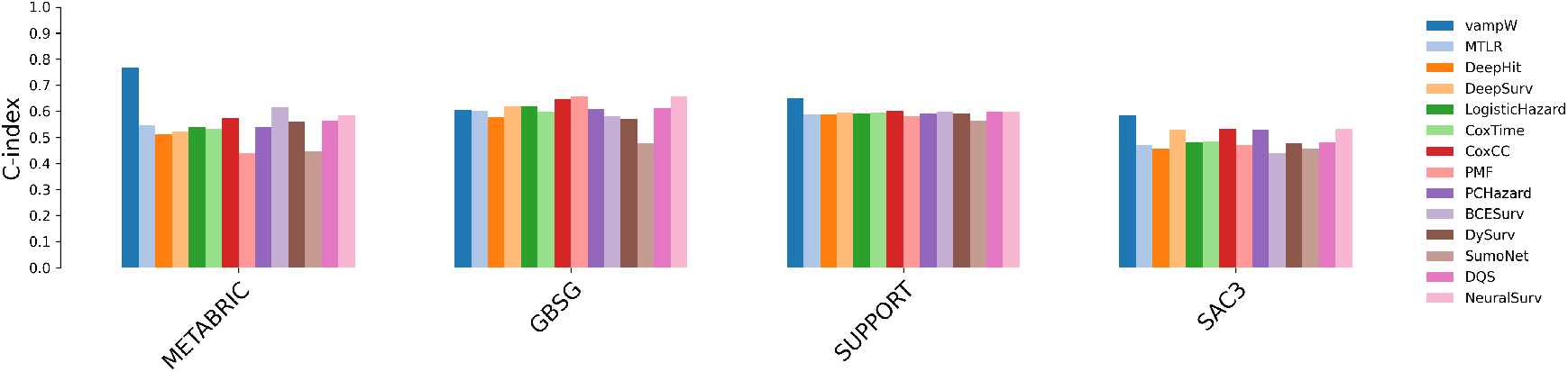
Out-of-sample performance comparison in terms of the Concordance index for vampW and deep learning methods. We compare predictions of vampW with common deep learning methods on several widely used survival benchmarking datasets, namely METABRIC, GBSG, SUPPORT, and SAC3. The benchmarking results for deep learning are reported in a recent NeuralSurv study [32], which addresses prediction in data-scarce regimes. vampW outperforms all deep learning methods on METABRIC, SUPPORT, and SAC3 datasets, and outperforms 6 out of 13 benchmarking methods on the GBSG dataset.

**Supplementary Figure 18:**
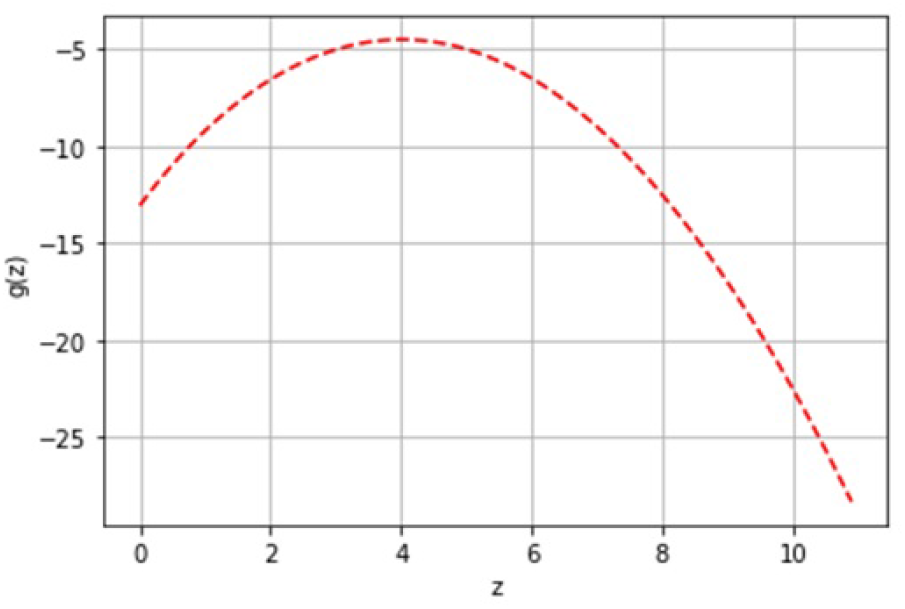
Visualization of a function *g*_*z*_ (*z*_*i*_) for *α* = 1, *τ*_1_ = 1, *p* = 5, *μ*_*i*_ = 0, *y*_*i*_ = 1.

### Supplementary Note

#### Details on denoiser for Weibull link function

We assume observations are independent, allowing for component-wise Maximum A Posteriori estimation. Proceeding with the construction of an age-at-onset denoiser for individual *i*, we express the posterior as follows, excluding terms independent of *z*_*i*_:

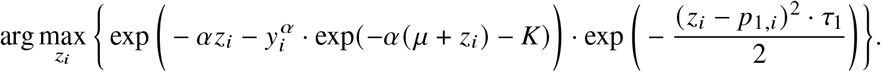

Let us define *g*_*z*_ : ℝ → ℝ as

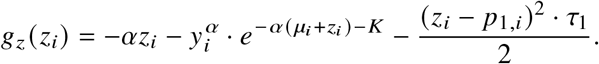

Then, taking the derivative with respect to *z*_*i*_, one obtains

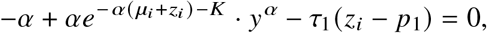

which can be solved using fixed-point iterations. In addition, we note that

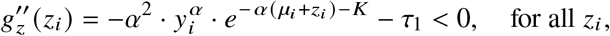

and, hence, *g* is a strictly concave function (see a visualization in Supplementary Figure 18).

Furthermore, by invoking the Implicit Function Theorem, we can compute the derivative of the denoiser as

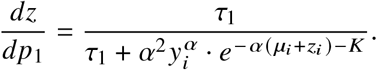

### Details on non-linear MMSE handling right-censoring

#### Lemma 1.

*Differentiating the function inside the optimization problem (7) with respect to* ***β*** *and equating the result to zero yields*

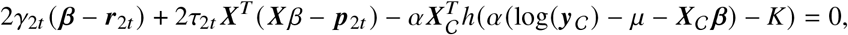

*where h* : ℝ → ℝ *is the standard hazard function of the Gumbel model given by h* (*x*) = *e*^*x*^, *and the notation h(****v****) with* ***v*** ∈ ℝ^*N*^ *denotes the component-wise application of the function h. Furthermore, the Onsager corrections for the denoiser and nonlinear MMSE step are equal to*

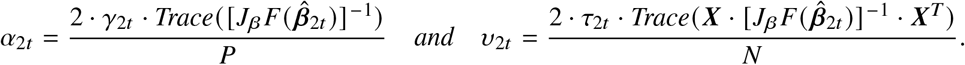

*Proof* Let us define the function *g* : ℝ^*P*^ → ℝ with

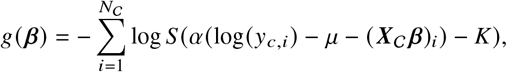

where *N*_C_ is the number of censored individuals in the study and ***X*** is the design matrix corresponding to those individuals. By performing differentiation, it follows that

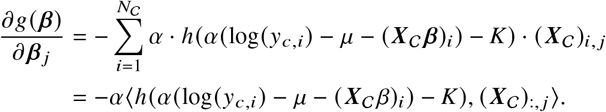

Hence,

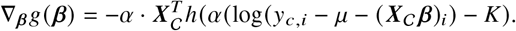

Furthermore,

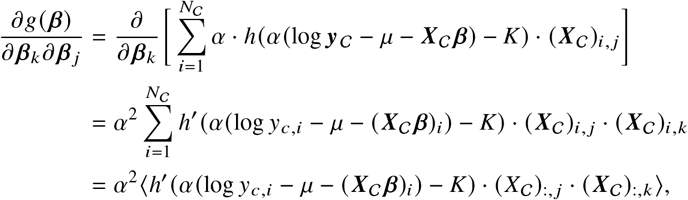

with *h*^′^ (*x*) = *e*^*x*^. It directly follows that

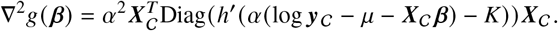

We now define *F* : ℝ^*P*^ → ℝ^*P*^ as

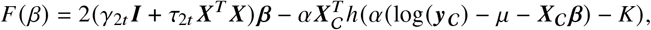

which means that a solution 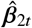 to the optimization problem (7) satisfies

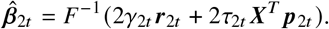

Note that

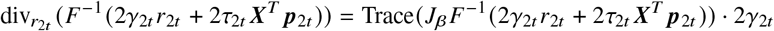

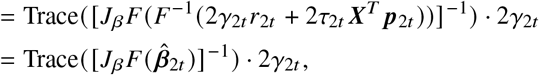

where

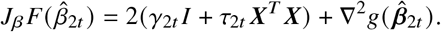

Finally, we conclude that

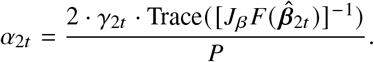

Similarly, we derive

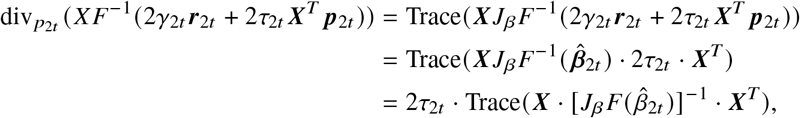

which gives the Onsager correction

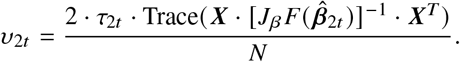

### Details on EM hyperparameter updates

We present the derivation of the expectation-maximization update formulae for the hyperparameters *μ* and *α* under the Weibull model for non-censored phenotypic outcomes. We define the E-step and M-step of the EM algorithm at iteration *t* of vampW in the following way:

1. <u>E-step:</u> The approximation of the posterior distribution typical for E-steps happens implicitly in VAMP-type algorithms, where the following approximate posterior probability distribution for the genetic component, ***z*** = ***X β***, is obtained:

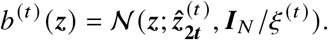
2. <u>M-step:</u> We aim to maximize the evidence likelihood over *μ* and *α*

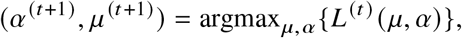

where we define

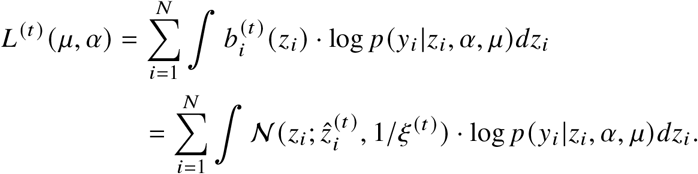

Here, 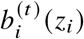 is the marginal distribution of the *i*-th component of ***z*** obtained from *b*^(*t*)^ (***z***) and; *N* (*μ, σ*^2^) denotes the probability density function of a Gaussian with mean *μ* and variance *σ*^2^ evaluated at *x*. Furthermore, note that

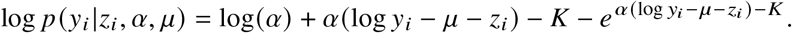

#### Update formula for *α*

By taking the derivative of *L* (*μ, α*) with respect to *α* and equating to zero we obtain:

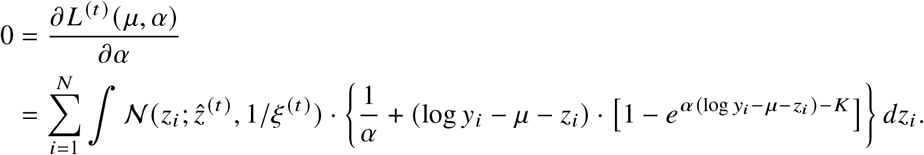

Separating the term in brackets results in

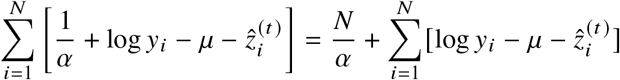

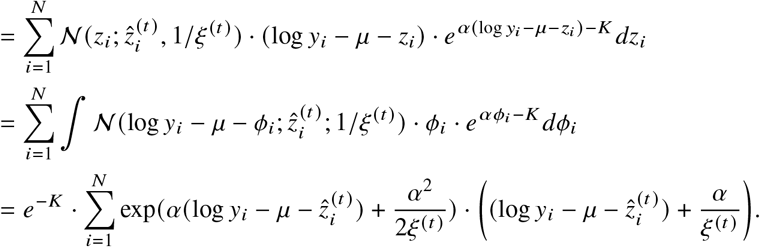

Finally, we define 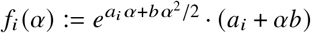, where 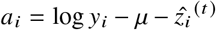 and 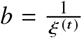.

It follows that

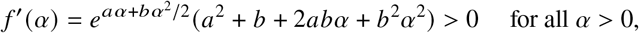

i.e. the function is strictly increasing in *α*. This implies that the first derivative with respect to *α* of the likelihood function *L*^(*t*)^ is

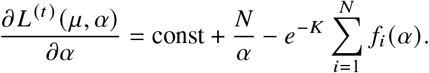

The second derivative is then given by

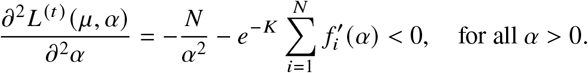

We thus have that the second derivative with respect to *α* is negative on the considered domain and hence the root of *∂*_*α*_*L*^(*t*)^ (*α, μ*)= 0 attains a unique global maximum on the considered domain. We solve for the global maximizer *α*^∗^ using numerical methods (SciPy optimize).

#### Update formula for *μ*

Similarly, by taking the partial derivative of *L*^(*t*)^ (*μ, α*) with respect to *μ* and equating it to zero, we obtain

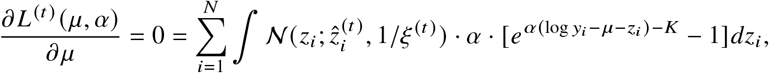

where we use that

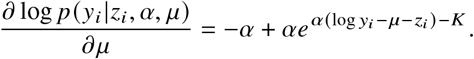

By separating the term in brackets it follows that

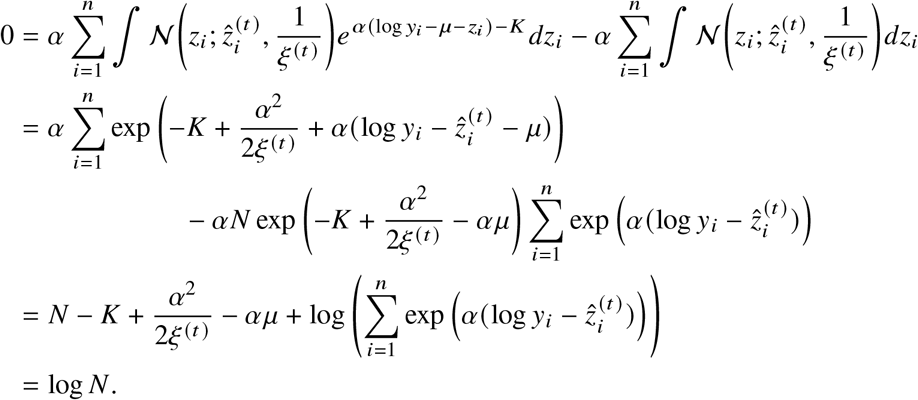

Hence, the update formula for *μ* satisfies

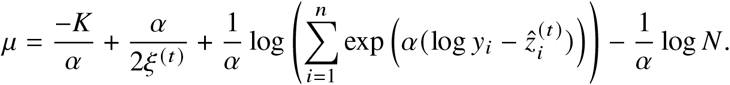

Furthermore, the maximization problem at hand is concave:

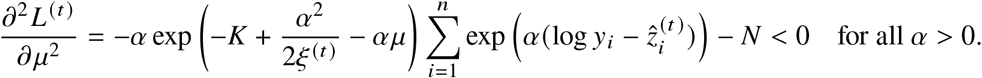

Therefore, *μ* satisfying 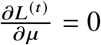 attains the global maximum of *L*^(*t*)^ (*μ*).

### Details on Weibull and ExpGamma outcomes

For the Weibull model, we calculate the scaling factor

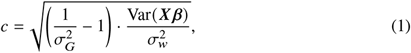

where 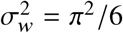 is the variance of the Gumbel distribution. Then, we generate time-to-event phenotypes as:

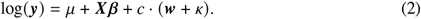

This formulation means that, in the phenotype log (***y***), the targeted proportion of variance is explained by markers.

For ExpGamma-distributed phenotypes, we exploit the fact that if *x* ExpGamma 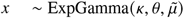, then 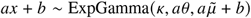. Then, we rewrite the model as follows:

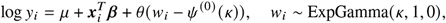

with shape parameter *κ*, scale parameter *θ*, and location parameter *μ*. We note that

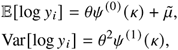

where *Ψ*^(*m*)^ (·) stands for the polygamma function of order *m*. Using the parametrization

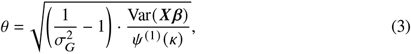

we obtain the time-to-event phenotypic outcomes, with the desired proportion of variance explained by the simulated markers.

## Footnotes

1 https://documentation.dnanexus.com/developer/api/running-analyses/instance-types

2 https://lifelines.readthedocs.io/en/latest/Examples.html

